# The Visual Dictionary of Antimicrobial Stewardship, Infection Control, and Institutional Surveillance

**DOI:** 10.1101/2021.05.19.444819

**Authors:** Julia Keizer, Christian F. Luz, Bhanu Sinha, Lisette van Gemert-Pijnen, Casper Albers, Nienke Beerlage-de Jong, Corinna Glasner

**Author notes:** Equal contribution (Julia Keizer, Christian F. Luz). Equal contribution (Nienke Beerlage-de Jong, Corinna Glasner).

## Abstract

**Objectives:** Data and data visualization are integral parts of (clinical) decision-making in general and stewardship (antimicrobial stewardship, infection control, and institutional surveillance) in particular. However, systematic research on the use of data visualization in stewardship is lacking. This study aimed at filling this gap by creating a visual dictionary of stewardship through an assessment of data visualization in stewardship research.

**Methods:** A random sample of 150 data visualizations from published research articles on stewardship were assessed. The visualization vocabulary (content) and design space (design elements) were combined to create a visual dictionary. Additionally, visualization errors, chart junk, and quality were assessed to identify problems in current visualizations and to provide improvement recommendations.

**Results:** Despite a heterogeneous use of data visualization, distinct combinations of graphical elements to reflect stewardship data were identified. In general, bar (n=54; 36.0%) and line charts (n=42; 28.1%) were preferred visualization types. Visualization problems comprised colour scheme mismatches, double y-axis, hidden data points through overlaps, and chart junk. Recommendations were derived that can help to clarify visual communication, improve colour use for grouping/stratifying, improve the display of magnitude, and match visualizations to scientific standards.

**Conclusions:** Results of this study can be used to guide data visualization creators in designing visualizations that fit the data and visual habits of the stewardship target audience. Additionally, the results can provide the basis to further expand the visual dictionary of stewardship towards more effective visualizations that improve data insights, knowledge, and clinical decision-making.

## Introduction

The amount of and reliance on data increases with the increase of scientific publications and information technologies in healthcare practice [1,2]. The increased complexity of big data raises various issues to be resolved by innovative big data analytics. This includes integrating, analysing and visualizing data to translate into meaningful information [3,4]. Translating raw data into meaningful information and communicating it to specific target groups is a challenge [1]. Without this translation and communication, researchers and practitioners cannot optimally use the information, so that the true value of the data remains hidden. Data visualization, here defined as the graphical representation of quantitative information, can facilitate the transformation, memorisation, and communication of data to understandable and actionable information. Data visualization also aids in the interpretation of increasingly large and complex datasets (big data) and in the understanding of sophisticated statistical models (machine learning) and their results – two rising trends over the last decades [5,6]. The importance of data visualization can, once again, be observed in the COVID-19 pandemic with the ubiquitous presence of charts, figures, and dashboards that aim to inform and support decision-making for a wide variety of target audiences [7].

Data visualization is a very active (research) field in itself and is generally part of typical statistical software used in the data analysis process (e.g. R, SPSS, SAS, STATA, Excel). Information and recommendations for the data visualization process are numerous and can be transferred between research fields or domains [8–11]. However, research on the visual domain context within a research field is often lacking, i.e. what the target audience is accustomed to see and expects in terms of content and design, and how this influences the perception and interpretation of data visualizations from different perspectives [12]. Common data visualization practices in a specific domain can be identified by studying the visualization design space [13]. This visual design space can be described as “an orthogonal combination of two aspects”, namely marks (i.e. graphical elements such as points, lines and areas) and visual channels to control their appearance (i.e. aesthetic properties such as colour, size and shape) [13].

To clarify the conceptual definitions for assessing and describing data visualizations a linguistic analogy can be used: a dictionary describes language in terms of both vocabulary (i.e. the set of words familiar in a language) and grammar/punctuation (i.e. the set of structural rules and supporting marks that control the composition and navigability of sentences, phrases, and words). Similarly, the visual dictionary describes visualizations in terms of both visual vocabulary (i.e. the domain content in terms of visualized data attributes) and visual design space (i.e. graphical elements and supporting aesthetic properties). The language or visual domain context is an overarching concept that represents language/visualization in practice, i.e. expectations and customs of the target audience, and how this affects their perception and interpretation of data visualizations (see also Figure 1). The visual domain context is, just as language, subject to changes over time and subject to interpretation differences based on varying perspectives.

**Figure 1.**
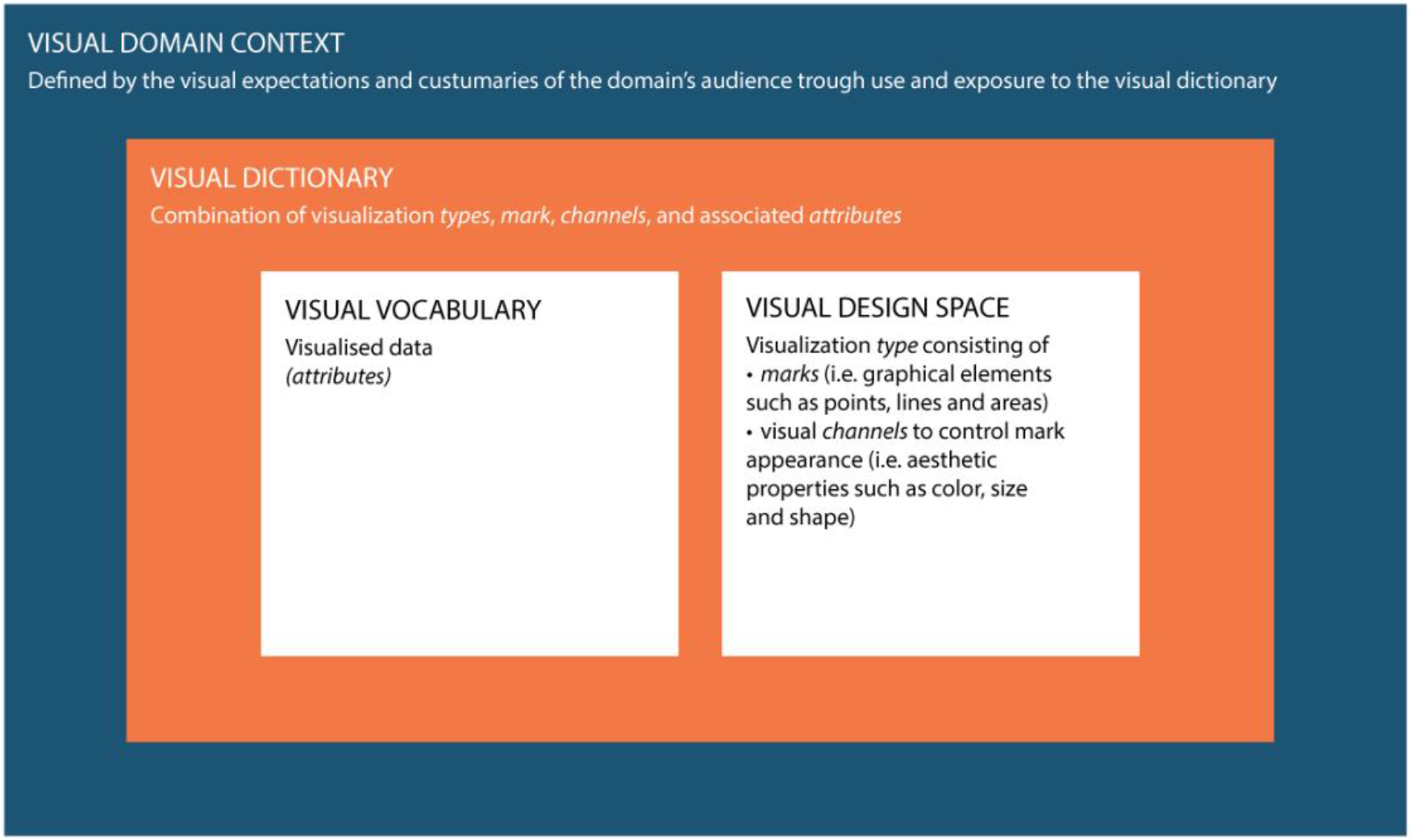
Conceptual framework used in this study to clarify the definitions and interrelations between the visual domain context, the visual dictionary and the visual domain vocabulary and visual design space.

Data and data visualization play important parts in the field of infectious diseases and antimicrobial resistance (AMR) for the reporting on the growing burden on health and healthcare systems [14,15]. Comprehensible and actionable information on antimicrobial consumption, pathogen distribution, or incidence and prevalence of (multi-) drug resistant microorganisms are vital to design interventions to tackle the AMR challenge [16]. On the hospital level, antimicrobial and diagnostic stewardship, infection control, and institutional surveillance (further summarised under ‘stewardship’) are the core components of strategies that promote the responsible use of antimicrobials and improve the quality and safety of patient care [17,18]. Data visualization is an integral part of these strategies, as it unveils the local situation and the drivers of AMR and can have a significant impact on the use of antimicrobials [19,20].

It has been shown how important it is to study data and data visualization experiences and perceptions in the medical domain and how this can influence the interpretation of data [21,22]. Depending on the type of visualization used, identifying the key messages from the data visualization can be substantially hindered. The audience’s background and its familiarity with data visualization (the visual domain context) have to be taken into account in the design process to avoid these obstacles. Example studies that identified the visual domain context by studying the design space can be found in the field of genomic epidemiology and genomic data visualization [23,24]. Although, some recommendations and best practices exist that are helpful in the data visualization creation process, common data visualizations practices in the field of stewardship have yet to be revealed [25,26]. The visual domain context and the use of data visualization in the field are unstudied – a systematic approach to define the design space is missing.

In this study, we aim to fill these gaps by assessing and defining the design space of data visualization in stewardship and to create a visual dictionary. The results of this study can help data visualization creators, such as healthcare/AMR/data professionals and scientists, to anticipate the visual domain context of the target audience and link it with existing recommendations for the data visualization process. This could benefit both research and clinical decision-making in the translation and communication of data to understandable and actionable information needed to tackle the AMR challenge, thereby improving the quality and safety of health and healthcare.

## Methods

### Study data

This study was based on a previous mapping study that clustered the field of AMR into 88 topics [27]. The map was generated by assessing the entire body of AMR literature available on PubMed between 1999 and 2018 consisting of 152780 articles. The identification of the 88 topics within the field was performed based on the title and abstract text using a machine learning algorithm (STM) [28]. The present study used all articles of three of the identified topics: *stewardship* (n = 3383 articles)*, infection control* (n = 1687 articles), and *institutional surveillance* (n = 2176 articles). Within the corpus of the 88 topics, these three topics reflect the core components of an integrated, comprehensive stewardship concept in institutional healthcare as defined above [18].

For each topic, a sample of 60 articles that contained at least one data visualization was randomly drawn. Data visualization was defined as the graphical representation of quantitative data. Geographical maps and flowcharts were excluded. From the sampled articles, one visualization per article was randomly sampled resulting in 180 data visualizations. The study design is shown in Figure 2.

**Figure 2.**
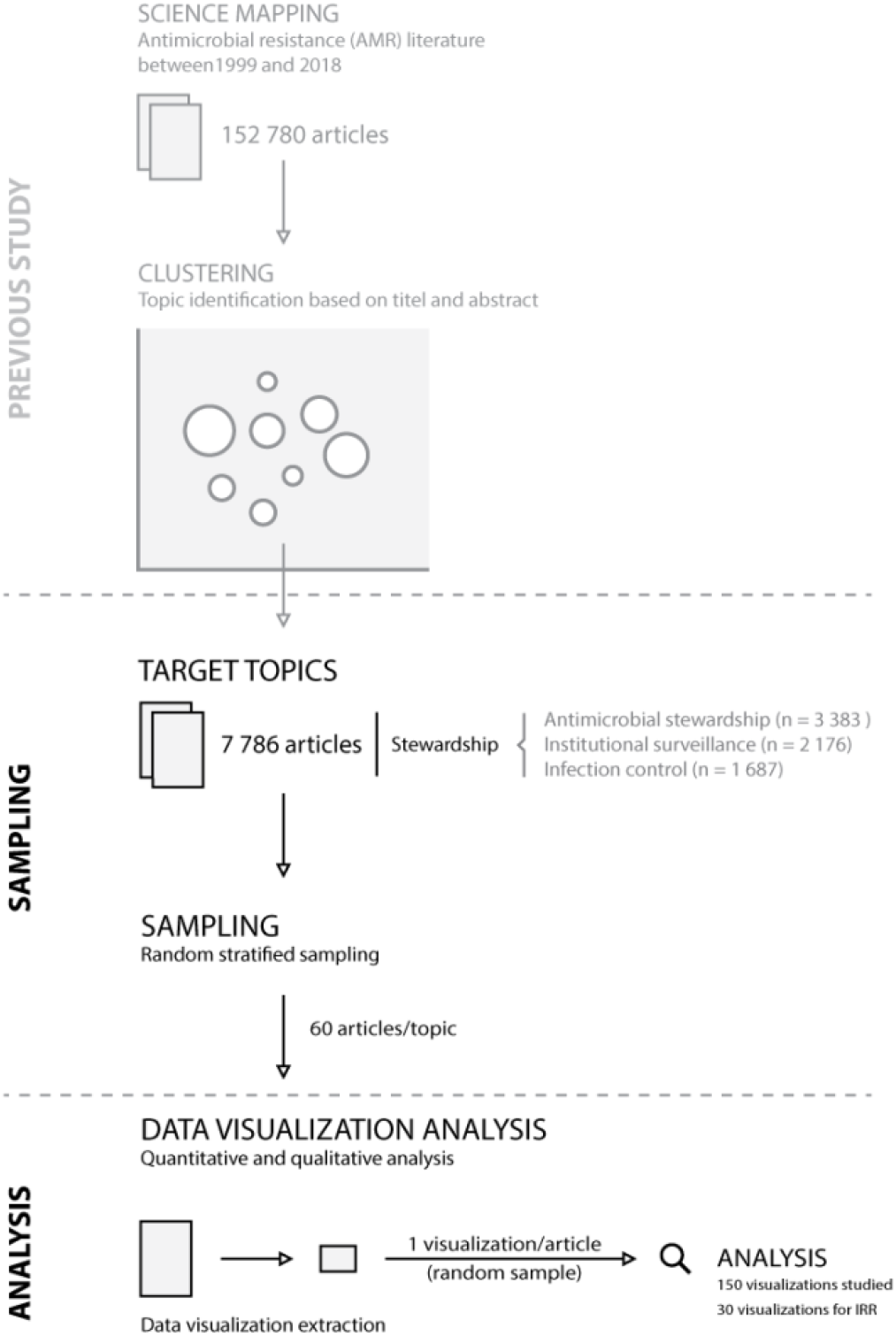
Study design. IRR = inter-rater reliability

**Figure 3.**
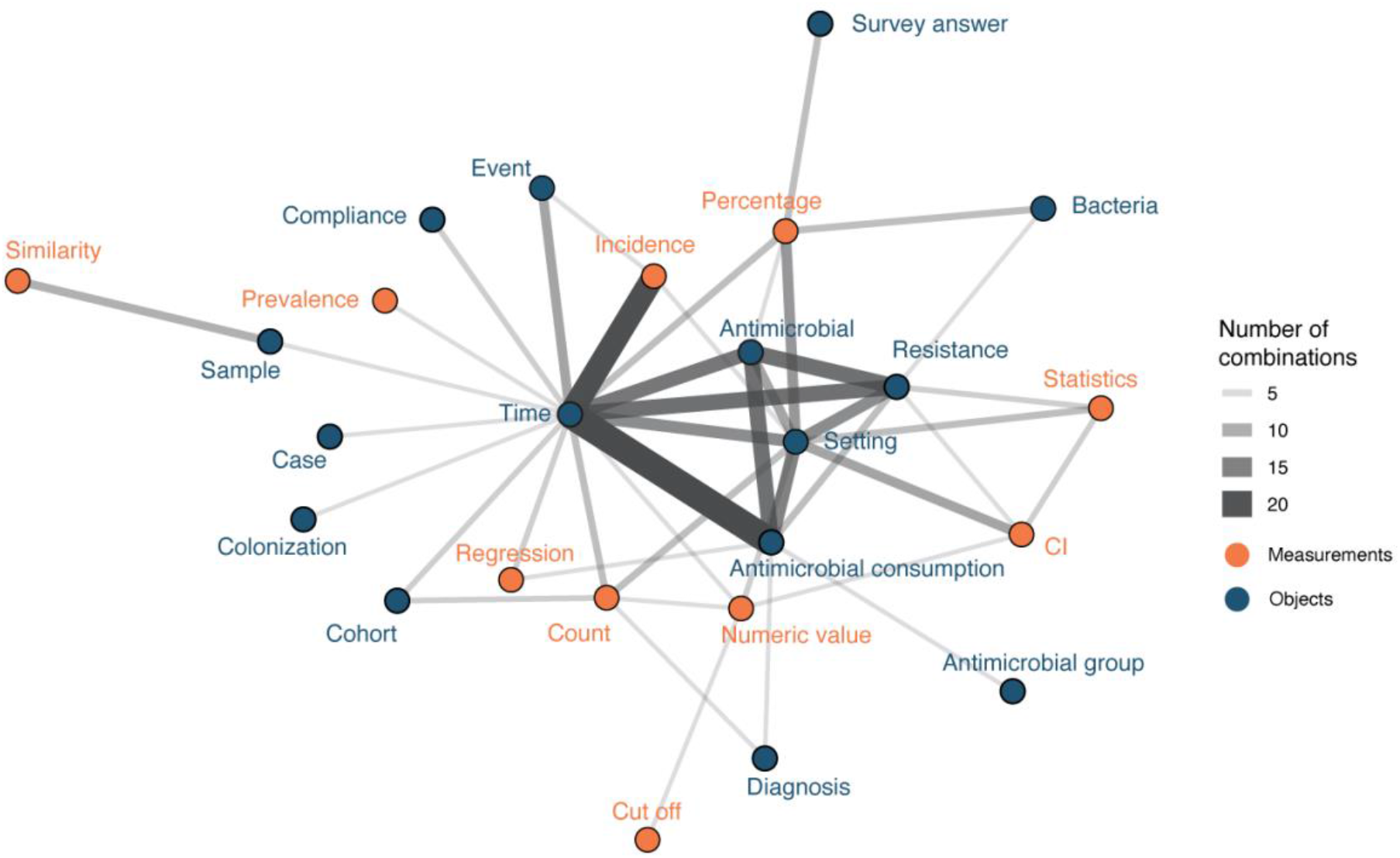
Attribute combinations in visualizations (combination count ≥ 3), thickness of lines corresponds to combination count. Orange points and labels represent attributes related to measurements; blue points and labels represent attributes related to objects.

To analyse reliability, ten randomly picked data visualizations of each topic were analysed in duplicate to calculate the inter-rater reliability (joint probability of agreement) [29]. Subsequently, 150 visualizations were included in the final analysis.

### Data visualization analysis

The extracted data visualizations were analysed based on the nomenclature and categorization developed by Munzner and further adapted for this study [13]. This approach dissected data visualizations into visual characteristics:

- *Attributes (or variables, parameters, features)*: the underlying data labelled as categorical, ordered, or quantitative.
- *Marks*: the basic geometric element (points, lines, or areas)
- *Channel:* channels control the visual appearance of *marks*

- *Position:* horizontal, vertical, both
- *Colour*
- Shape
- Tilt
- *Size:* length, area, volume

- *Channel effectiveness*

- *Magnitude:* ordered *attributes* can be expressed in ranks from most effective to least effective: position on common scale (most effective) > position on unaligned scale > length > tile/angle > area > depth > colour luminance/saturation > curvature/volume (least effective)
- *Identity:* the effectiveness to express categorical *attributes* can also be ordered: colour hue > shape

In addition, data visualizations were labelled with the visualization type used (e.g., bar chart, line chart, scatter plot, etc.) and the use of faceting (multiple linked visualizations in a design grid). Each visualization was assessed upon its interpretability without additional text (yes, if interpretable without additional information; partially, if a description was given in a caption; not all, if a description was absent or only available in the article text).

Visualization quality was captured by rating the first and last impression during the analysis process on a scale form 1 (poor) to 5 (good). The choice of the visualization type given the underlying data was rated on a scale from 1 (poor) to 5 (good). In addition, free, written text was recorded to capture comments and remarks about the data visualization.

A structured assessment form (supplementary materials S1) was developed comprising all the above mentioned elements.

The form was discussed within a multidisciplinary team of data-visualization and AMR experts. The assessment form was applied to each data visualization in a two reviewer (JK, CFL) process. First, the assessment form was used for training the analysis process with ten data visualizations not part of the final study data. Next, each reviewer analysed 50% of the study data visualizations followed by a re-review through the other researcher. Consensus was reached through discussion if the first assessment differed.

### Quantitative analysis

Results from the data visualization analysis step that were analysed with descriptive statistics comprised visualization type, number of attributes, faceting, rating, and visualization type choice. *Attributes* were analysed for pairwise co-occurrence and presented if a combination occurred more than twice in total.

### Visual dictionary

The visual dictionary was created based on the visual vocabulary content (stewardship-related data) and the visual design space (characteristics used to design the visualization). The content was analysed by identifying the *attributes* and grouping the *attribute* names using inductive coding. This part built the vocabulary of visualized stewardship data. Next, stratified quantitative analyses of visual characteristics (*channel, marks,* etc.) per *attribute* were performed, thereby adding the visual design space to the vocabulary to create the visual dictionary. Linking text and visual vocabulary enabled the creation of a visual dictionary to help identify *attributes* (e.g., resistance) with associated *channels* (e.g., points and lines on a common scale).

### Qualitative analysis

Comments and remarks about the data visualizations were coded in Microsoft Excel by two researchers (CL and JK). An open coding round was followed by axial coding to discover related concepts in the sub-codes. Differences were discussed until consensus was reached, which increased the internal validity [30]. Next to improvements, CL and JK coded remarks about chart junk (i.e. the unnecessary and/or redundant use of visualization embellishments) [11].

## Results

In total, 150 visualizations were analysed (IRR: 87% joint probability of agreement). The following sections are separated into visualization vocabulary (content) and visual dictionary with results stratified by identified *attributes*. These sections are followed by visualization ratings, identified visualization problems (including chart junk), and suggested recommendations for visualization creators and users.

### Visual vocabulary (content)

In total, 48 different coded attributes were identified. The majority (54.7%) of visualizations used three attributes. Two or four attributes were used in 18.0% and 20.7% of all visualizations, respectively. Few of the studied visualizations (6.7%) used more than 4 attributes.

The ten most used attributes were *time* (n=69, 46.0%), *setting* (n=43, 28.7%), *antimicrobial consumption* (n=32, 21.3%), *resistance* (n=31, 20.1%), *antimicrobials* (n=27, 18.0%), *percentage* (n=26, 17.3%), *count* (n=24, 16.0%), *incidence* (n=24, 16.0%), *numeric value* (n=20, 13.3%), and *bacteria* (n=12, 8.0%). Attributes can be grouped into objects (e.g. *bacteria*) and measurements (e.g. *percentage*). However, the following analysis focuses on attribute combinations and attributes are thus kept ungrouped.

Attributes showed different co-occurrence patterns (Figure 2). The ten most frequent combinations were *time* and *antimicrobial consumption* (n=21), *time* and *incidence* (n=18), *antimicrobial consumption* and *antimicrobials* (n=12), *antimicrobials* and *resistance* (n=12), *time* and *resistance* (n=12), *time* and *antimicrobials* (n=11), *antimicrobial consumption* and *setting* (n=10), *resistance* and *setting* (n=9), *time* and *setting* (n=9), and *percentage* and *setting* (n=8).

### Visual dictionary

#### Visualization types

Fourteen different visualization types were identified of which bar charts (n=54, 36.0%) and line charts (n=42, 28.1%) were predominantly used. Bar charts were most frequently associated with the attributes *antimicrobials, bacteria, cohorts, compliance, counts, diagnosis, errors, percentages, resistance, setting,* and *survey answers.* Line charts were predominantly associated with *antimicrobial consumption, costs, cut-off, incidence, numeric values, regression, statistics,* and *time* (detailed results available in the supplementary materials S2)

Different visualization types combined in one visualization were used in 10.7% (n = 16) of all visualizations. In these, visualization types that were combined more than once were bar charts with line charts (n=5, 31.3%) and stacked bar charts with line charts (n=2, 12.5%). In 41 visualization (27.3%) facets were used, i.e., one visualization split into a matrix of visualizations using the same axes.

#### Visual design space

Different patterns of visual characteristics could be identified for different *attributes* (detailed counts and percentages in supplementary materials S3).

1. Position: Horizontal axes were mostly used for *Antimicrobials, bacteria, confidence intervals, counts, cut-offs, diagnoses, events, numeric values, settings, similarity,* and *time*. In contrast, vertical axes were mostly used for *antimicrobial consumption, cases, cohorts, counts, errors, incidence, percentages, regression*, *resistance, samples, statistics,* and *survey answers.*
2. Marks, colour, shape: *Attributes* also differed in their use of marks. Some attributes had clear associations with mark types, e.g. *time* was visualised with lines in all instances. Areas as marks were seldomly used, e.g. for *antimicrobial consumption, counts, cut-offs, incidence, numeric values, percentages,* and *resistance.* Colour and shape channels were frequently used in most attributes. A detailed colour and shape channel analysis is available in the supplementary materials S3.
3. Size: Size was most often visually reflected through length. Area to reflect size was used for *antimicrobial consumption, count, cut-off, incidence, numeric values, percentages,* and *resistance.* Volume was rarely used (*count, percentages*).
4. Ordering: Position on a common scale was mostly used in quantitative and ordered attributes reflecting the best channel effectiveness for these attribute types. Categorical attributes mostly used colour hue, which is preferred over the less effective use of shapes. A detailed channel effectiveness analysis is available in the supplementary materials S4.

### Ratings, problems, and chart junk

#### Visualization ratings

Overall, 55.3% (n=83) of all visualizations were interpretable without additional text (in caption or in the manuscript text). The overall choice of visualization type for the presented data was rated with a mean of 4.62 (SD: 0.9) on a scale from 1 (poor) to 5 (good). The general assessment of the visualization quality (scale 1=poor to 5=good) was rated with a mean of 3.6 (SD:1.2). Identified problems (and recommendations) are described below.

#### Identified problems

The coding of the identified problems are presented in the coding scheme in Table 1, including axial codes, open codes and frequencies. The axial and open codes are further elaborated upon below the table. In supplementary materials S5, additional illustrative quotes per code are presented.

**Table 1.**
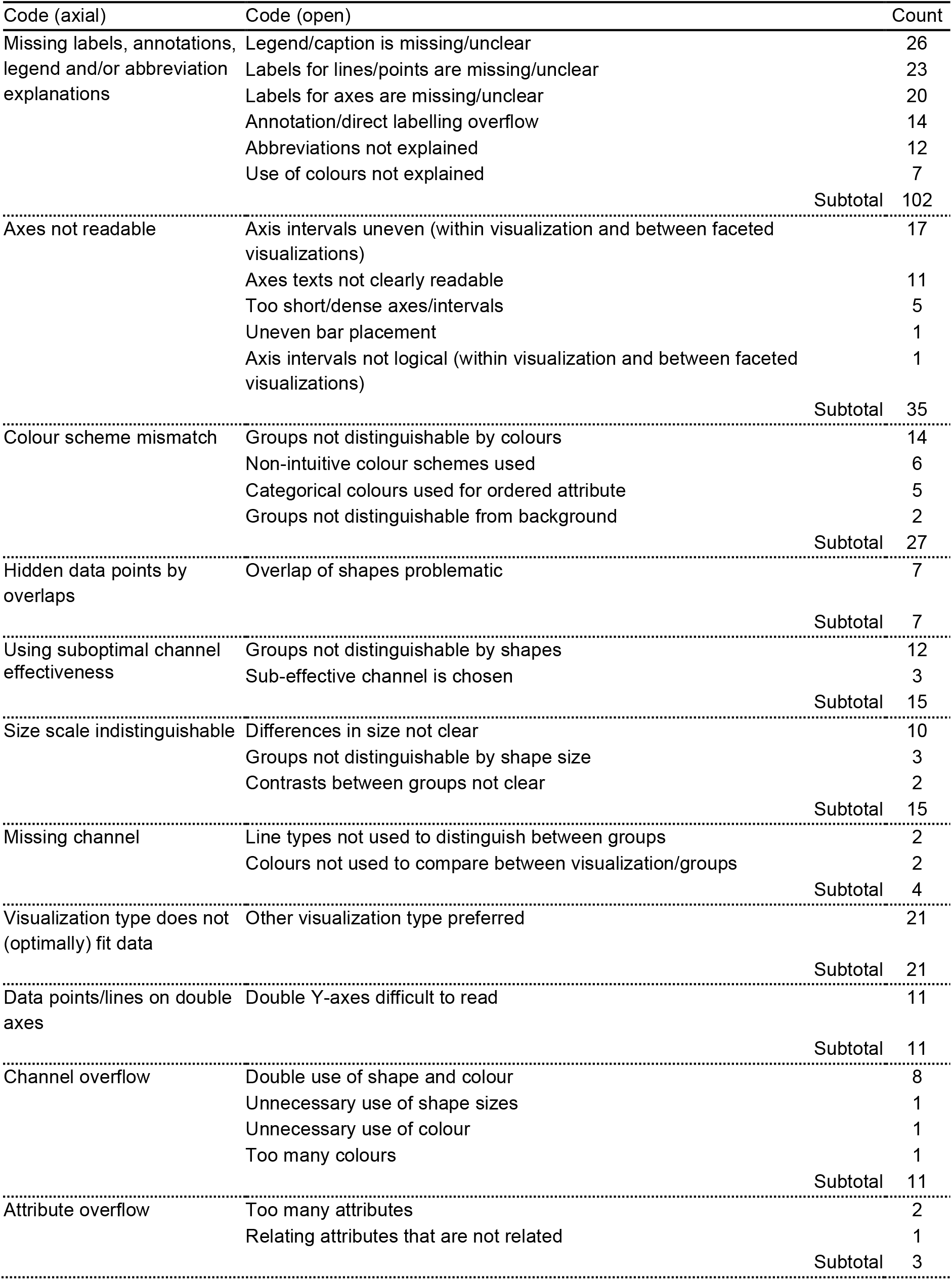
Identified problems in data visualization.

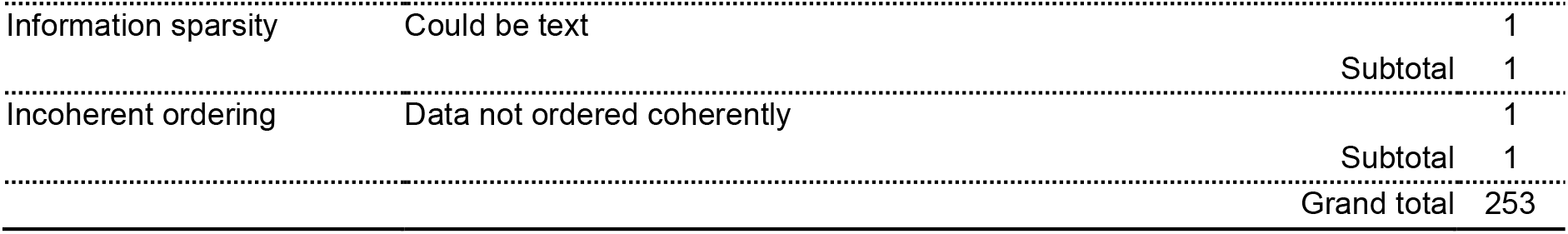

Most problems were related to the clarification of the visualization, because of missing or unclear labels, annotations, legend, captions and/or abbreviation explanations (n=102; 68.0%). Other problems concerned the axis readability (n=35; 23.3%) for example due to uneven axis-intervals within a visualization and between faceted visualizations for comparison. Problems related to distinguishing data points and groups in visualizations were detected, for example with mismatches in colour scheme (n=27; 18.0%), hidden data points by overlaps (n=7; 4.7%), and using suboptimal channel effectiveness (n=7, 4.7%). In some cases, data points and groups were not clearly distinguishable because of the size scale (n=15; 10.0%) or because of missing channels (e.g. line type or colour, n=4; 2.7%). Furthermore, problems identified were the suboptimal or wrong choice in type of visualization (n=21; 14.0%) and the confusing use of double y-axis (n=11; 7.3%). Some visualizations were overcrowded, either in terms of channel overflow (e.g. using both colour and shape, n=11; 7.3%) or attribute overflow (e.g. too many attributes, n=3; 2.0%). On the contrary, the information in one visualization was sparse enough to be written in text (i.e. no added value of a visualization). Lastly, one problem related to the incoherent ordering of data.

#### Chart junk

Most chart junk represented text that cluttered the visualization (n=8), for example with redundant direct labels for each data point. Other chart junk was found in visualizations using unnecessary 3D (n=8), background colours (n=6), shadow (n=4), and colour/shape filling (n=4).

#### Examples and recommendations

To illustrate problems in data visualization, we designed a visualization that exhibits several of the identified problems based on simulated data (Figure 4). Figure 5 proposes an alternative to Figure 4 where the identified problems were avoided. Of note, data such as the simulated data in these figures can be visualised in many different ways, depending on the underlying research questions.

**Figure 4.**
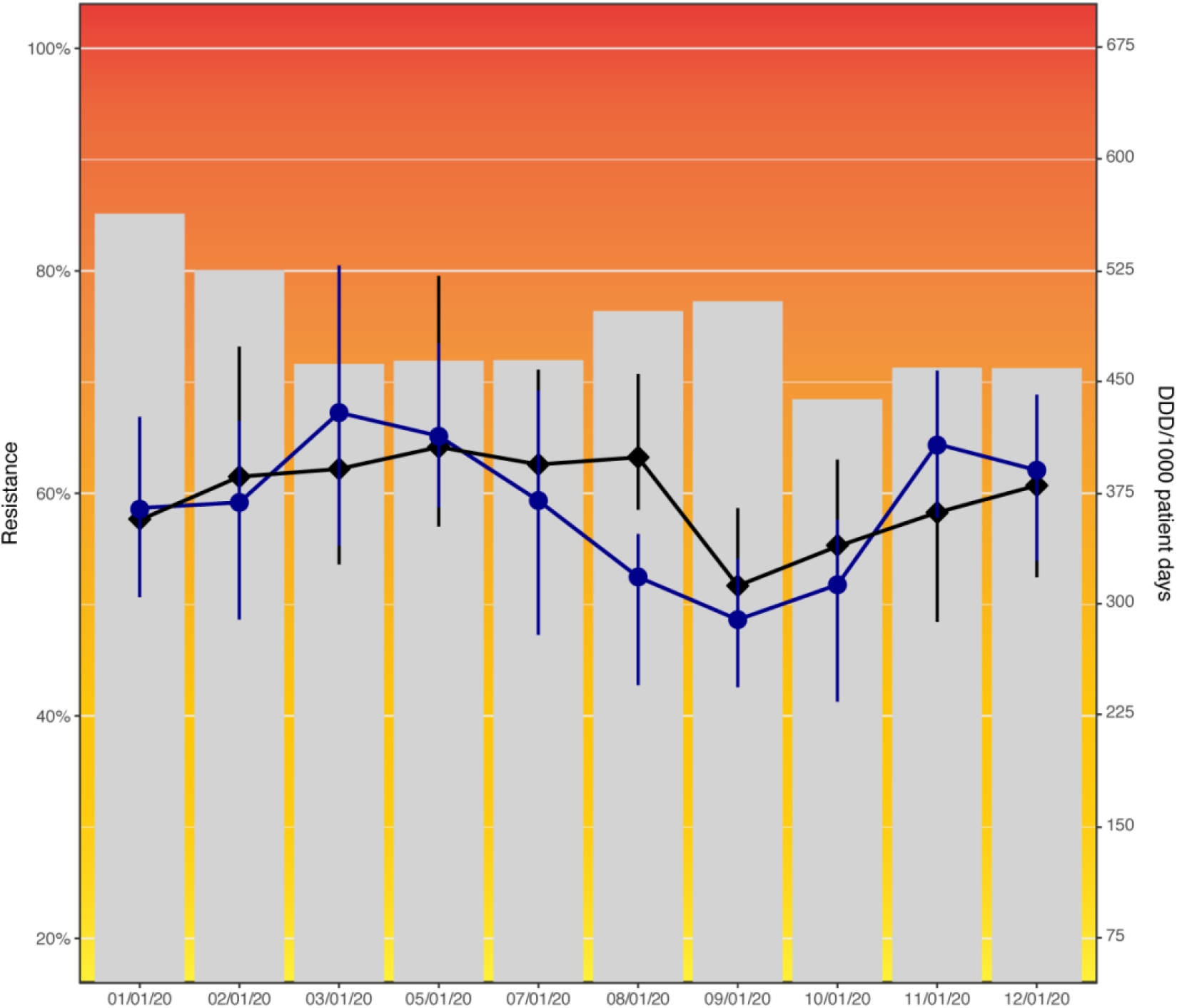
Resistance to amoxicillin in *Escherichia coli* and consumption of cefuroxime (black) and piperacillin/tazobactam (blue) across hospital departments in 2020. This data visualization (simulated data) shows several problems identified in this study: Axes not starting at zero, use of double y-axes, background colours, hidden data points by overlaps, colour scheme mismatch (blue and black difficult to distinguish), unequal axis steps on x-axis, missing legend, incomplete axis labels (abbreviation not explained).

**Figure 5.**
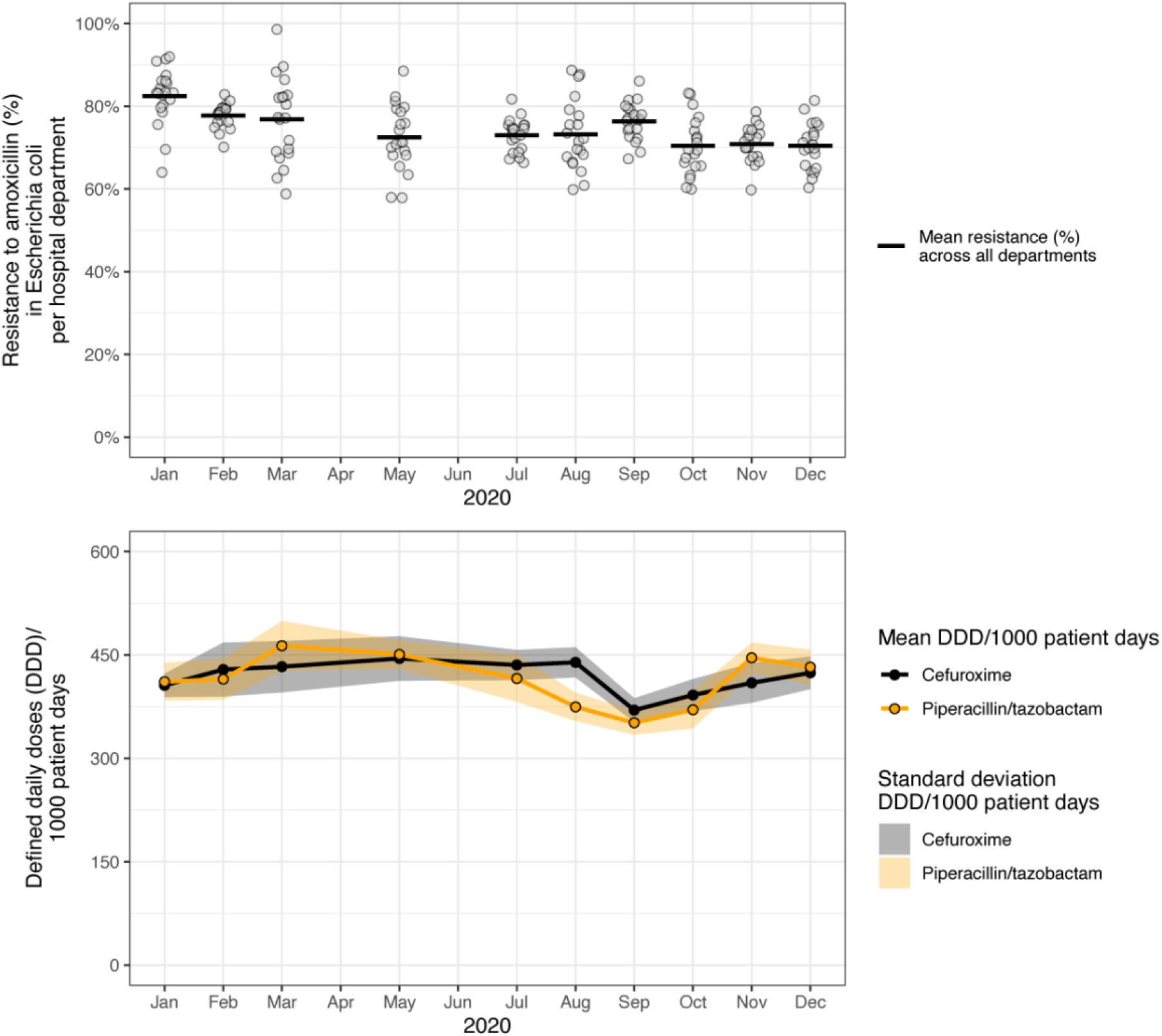
Resistance to amoxicillin in *Escherichia coli* and consumption of cefuroxime and piperacillin/tazobactam across hospital departments in 2020. These data visualizations use the same data as in Figure 4 (simulated data), but propose an improved visualization.

Figure 6 summarises the results of this study and presents the visual dictionary of stewardship. In addition, it provides a set of recommendations to avoid the most common problems in data visualizations as identified in this study.

**Figure 6.**
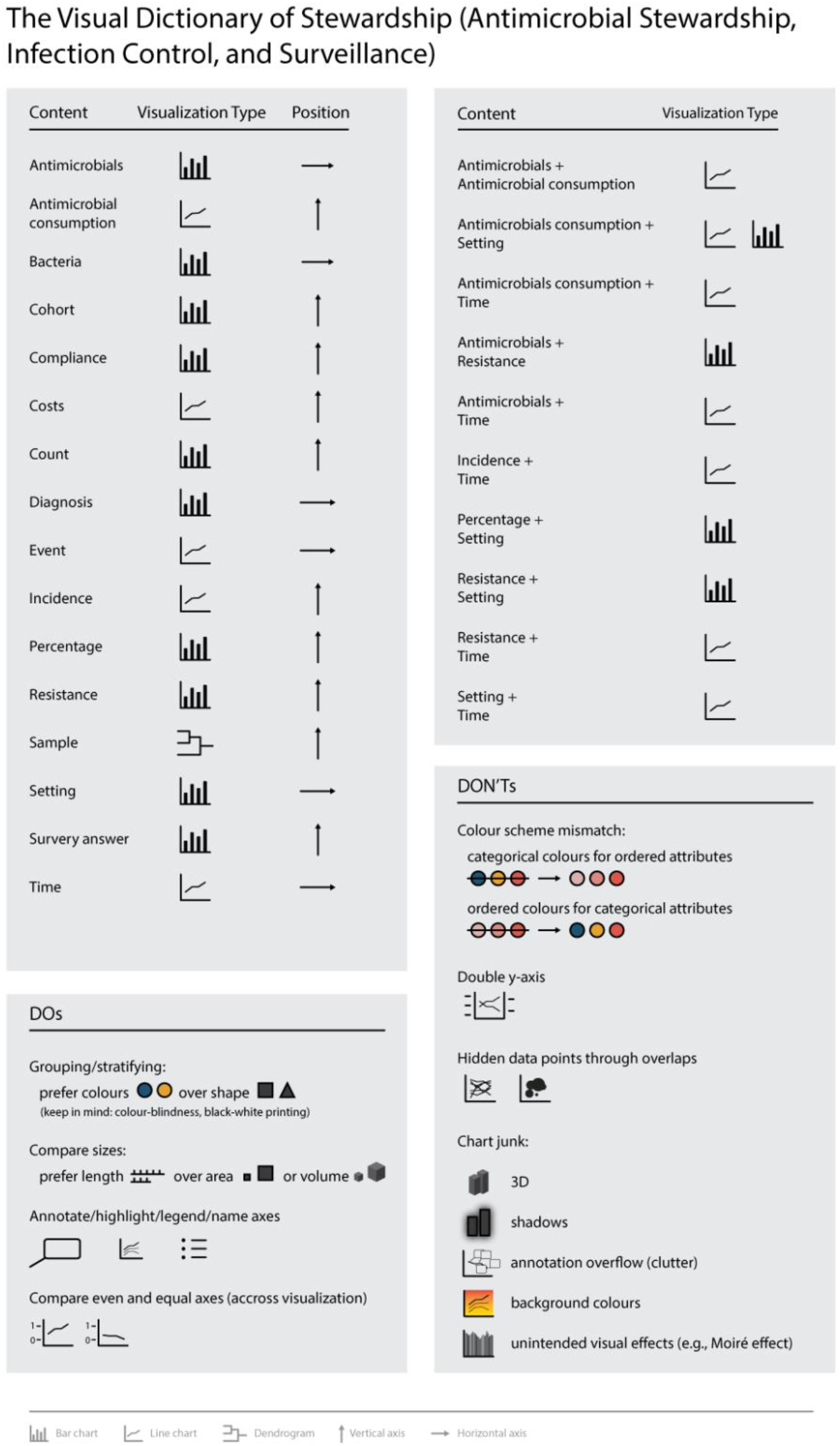
The visual dictionary of stewardship (antimicrobial stewardship, infection control, and institutional surveillance).

## Discussion

This study systematically analysed the visual domain context of stewardship (i.e. antimicrobial stewardship, infection control, and institutional surveillance). Stewardship healthcare experts and scientists that create data visualizations can benefit from the revealed visual domain context, since it allows them to anticipate the visual habits of their target audience. The results of this study can serve as the basis to inform visualization creators to optimise visual communication in the field and to guide user-centred design, e.g., in clinical decision support systems.

### Implications for data visualization creators

With the systematic analysis of the visual domain context of Stewardship we revealed common practices and identified problems with current visualizations. In this study we identified 14 different visualization types used in the visual domain context of the field. However, more than 80% of all visualizations used classical (stacked) bar or line charts; quite homogenous design choices. We argue that the visualization type choice is based on tradition and habits as a systematic approach to data visualization in the field was missing until now [26]. For researchers in the field that communicate their findings to various stakeholders (e.g. stewardship professionals, policy makers, epidemiologists) the described visual domain context in this study provides guidance to match their visualizations with the audience’s visual expectations and habits. Especially the visual dictionary, the link between often used attributes (i.e. content) and associated design choices (e.g. visualization type or marks), will help to compose visualizations that fit with common practice. However, given the wide variety of data in the field and the increased complexity that big data will add (in terms of volume, velocity, variety, veracity, validity, volatility and value), more “visual variability” might be expected and even needed in the future [3,31,32]. Informing and teaching visualization creators and users about data visualization design alternatives is an important step in this process. A lack of awareness and knowledge about data visualization design alternatives might lead to suboptimal data visualizations. Examples from our findings were the use of less effective visual channels, suboptimal plot types for the presented data, or mismatches in colour choices for different data types. These are examples of instances where the respective data visualization creators require more support in visualization design choices. We see a clear role here for data visualization experts and software developers to cocreate open-source/access tools that support visualization creators in their visualization choices (e.g. reminders for adding labels and legends, suggestions for optimal colour schemes, warnings in case of chart junk). Our results and findings from similar studies in other fields [23,24] can support them in doing so by providing an overview of what is already used, including potential pitfalls. Of note, academic journals play an important part in this process by providing the platform for data visualizations and should be encouraged to promote high quality data visualization practices.

### Improving data visualization practices

In general, the use of data visualizations for communicating data is highly encouraged. It greatly supports the interpretation, memorisation, and communication of insights and knowledge gained from data. Based on the identified problems with data visualizations in this study, several recommendations can be made to improve data visualization practices both in general and for stewardship specifically. Some recommendations relating to identified problems were already depicted in Figure 5 and 6 and are elaborated and extended upon below.

#### Colours

The use of colours in data visualization is highly complex. Colours can make a plot more appealing. Colours are also more effective than shape to distinguish categorical data [13]. Yet, shapes for categorical data were still widely used in the studied visualizations. This could reflect the need to provide visualizations that are black-and-white compatible (printable), although this has become less important with most of the scientific content being accessible online. Several aspects are key to consider when designing data visualizations with colour: colour-blindness, distortion through uneven colour gradients, or the perceived order of colours [33]. While field-specific colour codes might exist (e.g., red colour to represent resistance), general recommendations for the use of colours in scientific publications are available and are applicable across fields [33]. Extensive information on the use of colours in data visualization can also be found in online blogs from designers in the data visualization community (e.g., https://blog.datawrapper.de/which-colour-scale-to-use-in-data-vis/).

#### Adding statistics

Common scientific visualization types such as heatmaps or boxplots were rarely used in the studied data visualizations. In general, statistical aggregate parameters were often lacking. This would often have improved the visualizations under study. Boxplots are a classical example. However, this visualization type is rightfully criticised to conceal individual data points and could be misleading [34]. We also identified difficulties in displaying individual data points as one of the main problems. This problem was caused by overlaps, problematic scale sizes, or missing channels to distinguish data points. We highly advise to stick to the mantra of “above all show the data”, to avoid overlapping or concealing graphical elements [11], and to carefully balance the number of visualised attributes per visualization with simplicity.

#### Standardizing the visual dictionary

Although a large variety of data were displayed in the studied visualizations (48 different *attributes* such as antimicrobials, bacteria, or time), we observed some prominent patterns in the content and purpose of data visualization. Changes over time, e.g. time series, were part of 43.3% of all studied visualization. Twenty percent and 19.3% included antimicrobial consumption or resistance, respectively. This is not surprising given the importance of these data in the field. It could be worth considering to standardise data visualizations for these data types and contents similar to the consensus of international guidelines committees in the field, e.g. the European Committee of Antimicrobial Susceptibility Testing (EUCAST) or the Clinical & Laboratory Standards Institute (CLSI). This could help ensure high quality data visualizations for reliable insights in AMR- and stewardship-related data. Such initiatives to standardise data visualizations have already been taken by bodies in other fields, e.g. the Intergovernmental Panel on Climate Change (IPCC) [35]. Another example is the development and evaluation of a standardised medical data visualization method based on the ISO13606 data model [36].

### Limitations and strengths

This study has several limitations. Despite sampling from a comprehensive set of articles that cover the field of stewardship (antimicrobial stewardship, infection control, and institutional surveillance), only a limited sample of data visualizations were included. Data visualizations for content (attributes) not covered in this study could have been missed. However, the homogeneity of the identified data visualization types suggests that saturation was reached regarding the visual design space in the field. Another limitation is that we included data visualizations from scientific publications and not from other sources relevant to stewardship data visualizers (e.g. data systems used in practice [12,37] and AMR policy reports [38,39]). As a result, our findings might be more applicable to stewardship researchers and data visualization experts than healthcare professionals. Another use of data visualization, namely the exploration of increasingly complex big data, was outside the scope of the included articles [40,41]. Subsequent research into the visual domain context of stewardship should include these additional data visualization sources and applications to ensure a more comprehensive picture for healthcare professionals. Even though the extracted data visualizations were systematically analysed using a structured assessment form based on existing data visualization nomenclature and categorization [13], the analyses relied on the subjective interpretation and rating by the coding researchers. Several measures were taken to validate our findings, including discussing the assessment form and results within a multidisciplinary team of data-visualization and AMR experts, analysing the interrater-reliability, and comparing our findings to other data visualization studies. Our study is one of the first empirical studies that explores the use of data visualization in stewardship, thereby adding to the few review studies providing primers for data visualization recommendations and best practices in the field of antimicrobial stewardship, infection control, and institutional surveillance [25,26].

### Future perspectives

Future research can build upon our results by studying and expanding the use and design of data visualizations beyond the basic visual dictionary provided here. Two important future research directions are elaborated upon below.

Studying the visual domain context is as important as studying data visualizations themselves. This includes studying expectations and customs of the target audience, how this affects their perception and interpretation of data visualizations, and how this consequently impacts their decision-making or behaviour. The importance of assessing visual habits and perceptions in data visualization has been demonstrated before in other medical fields [21,42]. It was shown that personal preferences and the familiarity of a target audience with certain visualization types can result in tensions with data visualization recommendations and novel data visualization approaches. An example is the use of pie charts, which is often discouraged in the data visualization community. The target audience might still favour this type of visualization because of its apparent simplicity, despite the fact that pie charts are less accurately interpreted as angles and wedges are difficult to compare [8,43]. We strongly believe that incorporating best practices and data visualization recommendations are essential but advocate that these should be carefully balanced with visual habits and expectations in the field, and the message to be transported. An exemplary study is published by Aung *et al.* focusing on data visualization interpretation capacity and preferences in their target audience by combining interviews on interpretability and card-sorting of preferred visualizations [21]. Additionally, research is needed to better understand how data visualizations impact the viewers/users in terms of changes in opinions or attitudes that direct decision-making or behaviour changes [44].

In future research special attention should be paid to matching the visual dictionary and the context in which the visualization will be used in terms of users and their tasks and current practices (e.g. studying questions like “What kinds of visualizations are currently used?” and “How do they support to do current tasks?”) [45]. We see a clear parallel with user-centred eHealth design that emphasises the need of a holistic understanding of the interrelations between technology, people and their context [46]. Both qualitative (e.g. interviews) and quantitative (e.g. eye-tracking in current data visualizations) study designs can contribute to such a holistic understanding, which in turn can inform or improve the design of visualizations (or eHealth) in terms of required content, functionalities and usability [47]. Therefore, complementing research on data visualizations, as the current study and many other studies do, with research that primarily focuses on the interaction between people, their context and how data visualizations can support them, is needed [45].

## Conclusion

In this study, we analysed the visual domain context of stewardship (antimicrobial stewardship, infection control, and institutional surveillance). We successfully created a visual dictionary that can support the process of creating and using tailor-made data visualizations in the field. Thereby, our results allow data visualization creators to learn the *visual language* of the diverse field of stewardship. As data-driven solutions for stewardship are of increasing importance, effective processes of transforming this data to insights and knowledge is essential. Data visualization supports and enables this transformation and our results can guide the optimal visualization design choices that are grounded on expectations and habits in the field. In the future, our study can provide the basis to further expand the visual dictionary of antimicrobial stewardship towards more effective data visualizations that improve data insights, knowledge, and decision-making.

## Acknowledgement

The authors would like to thank Anamaria Crisan, Tamara Munzner, and Nils Gehlenborg for the inspiration provided for this study.

## Supplementary materials

### S1. Data Visualization Assessment Form

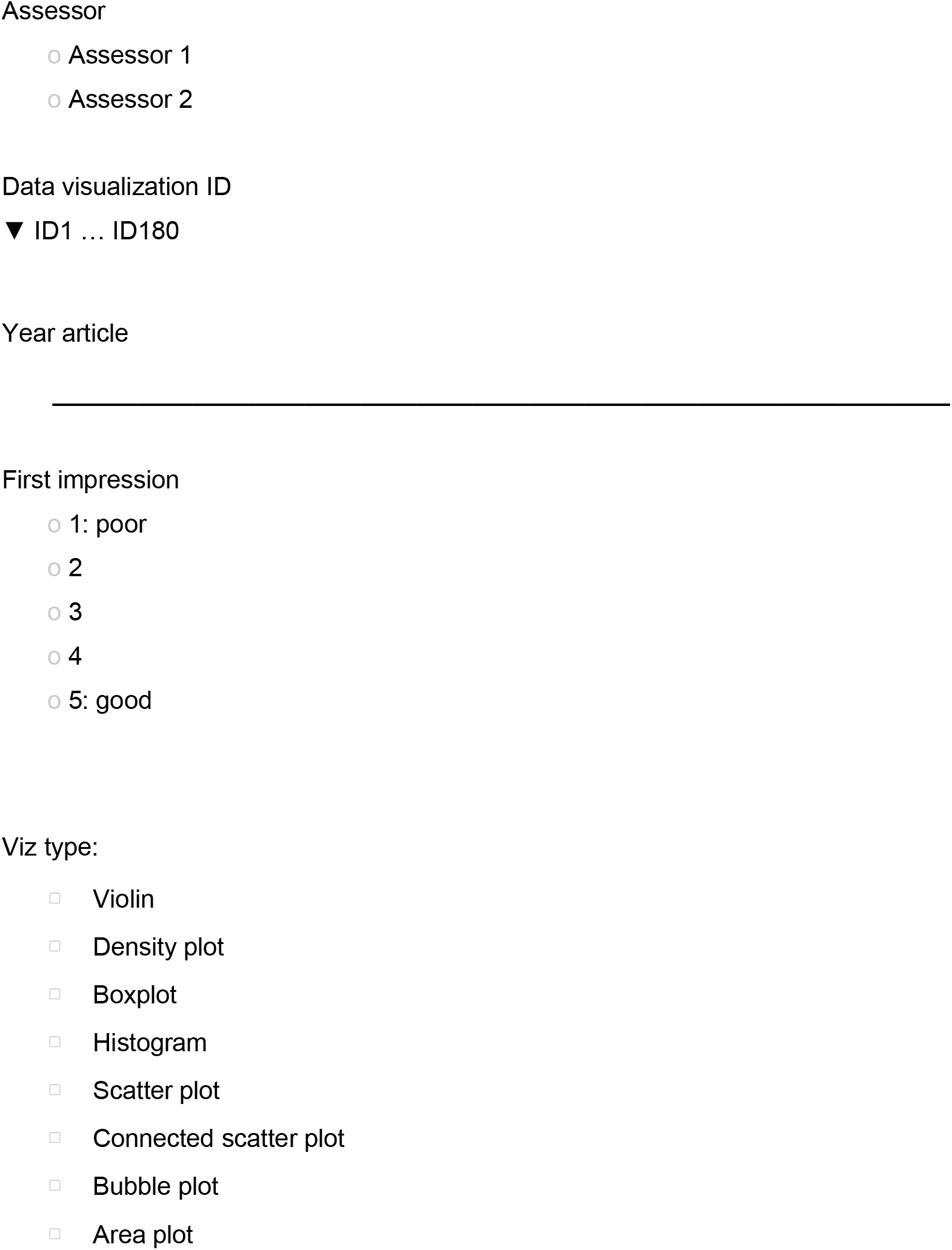

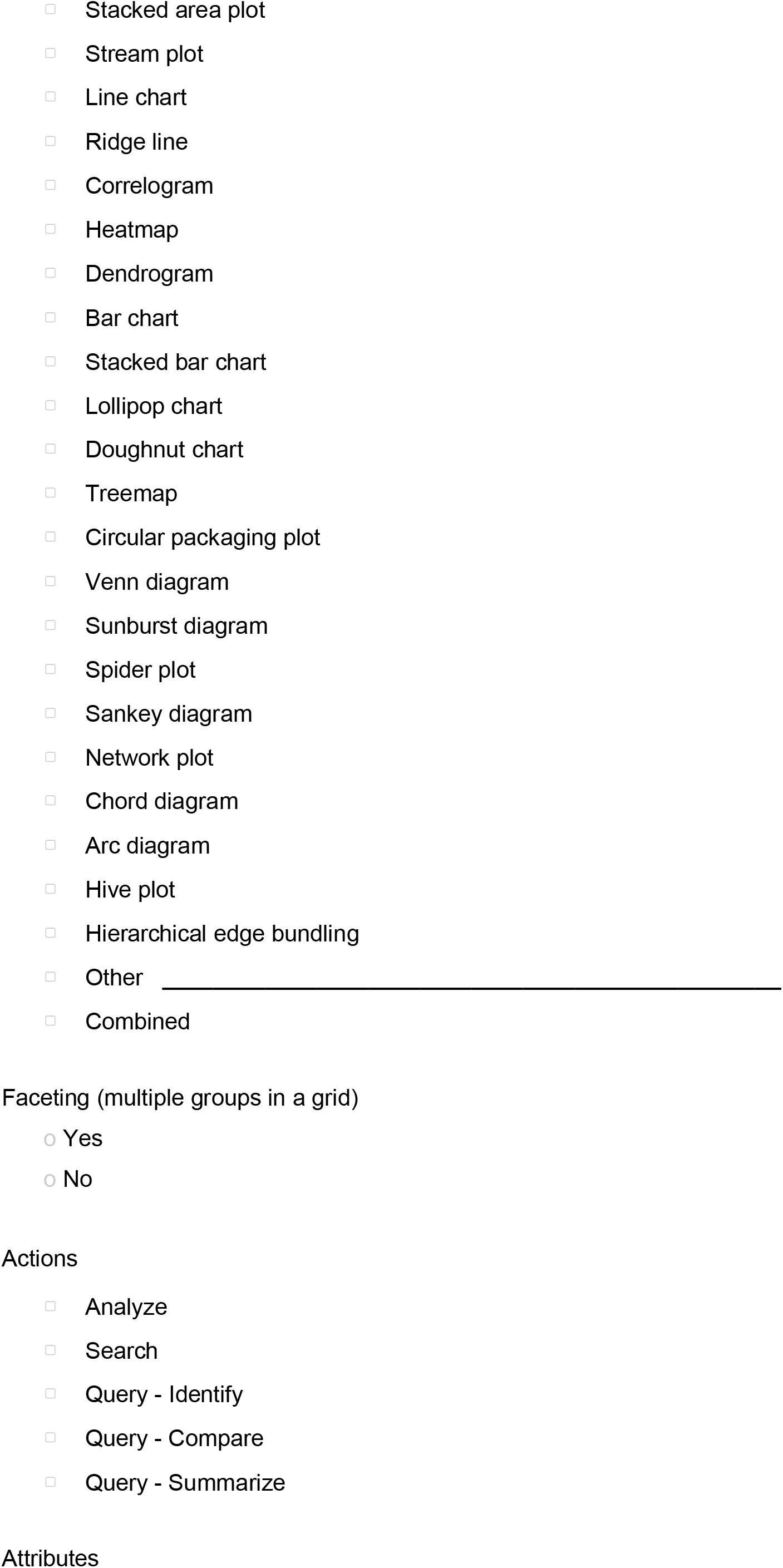

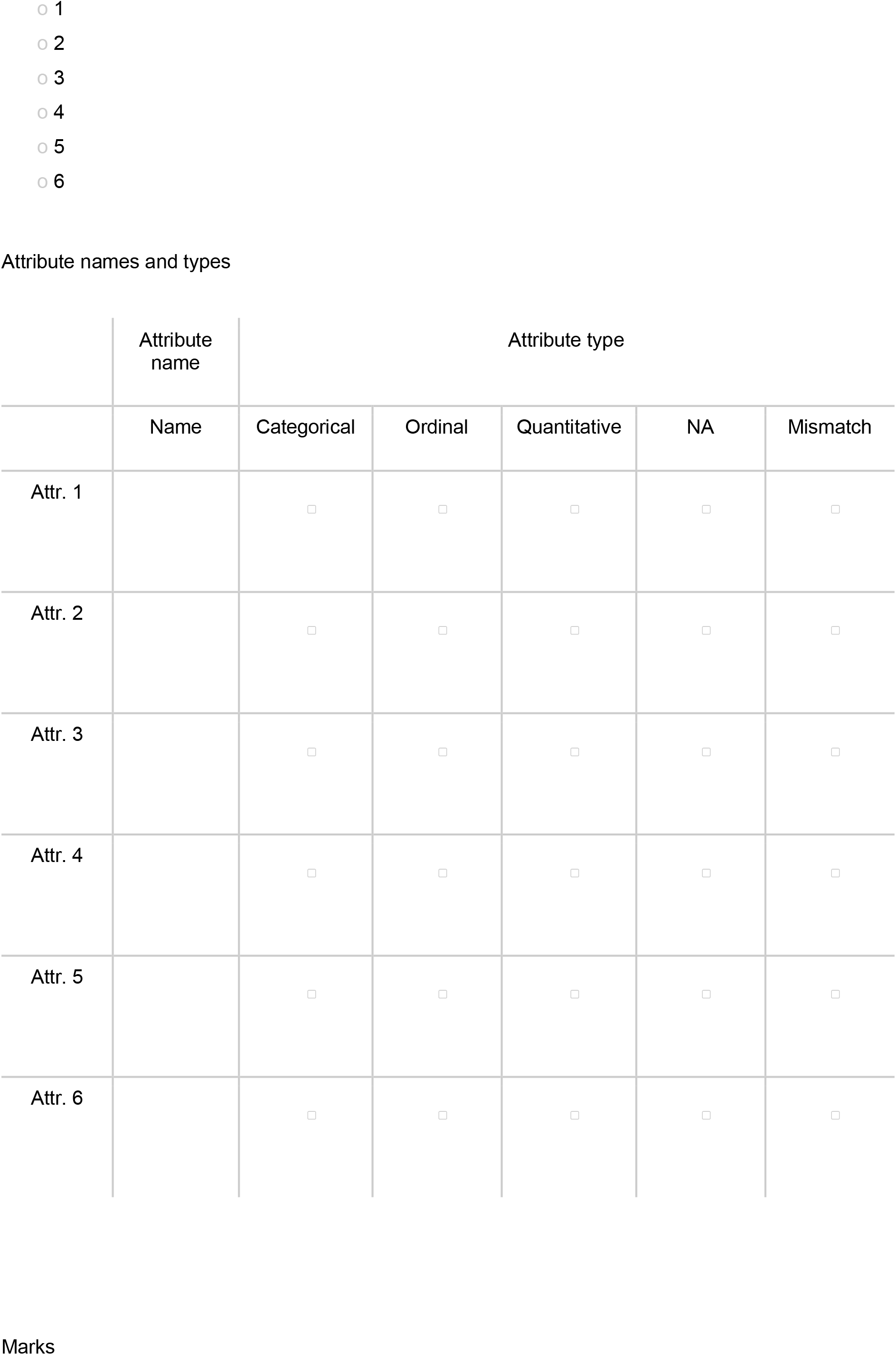

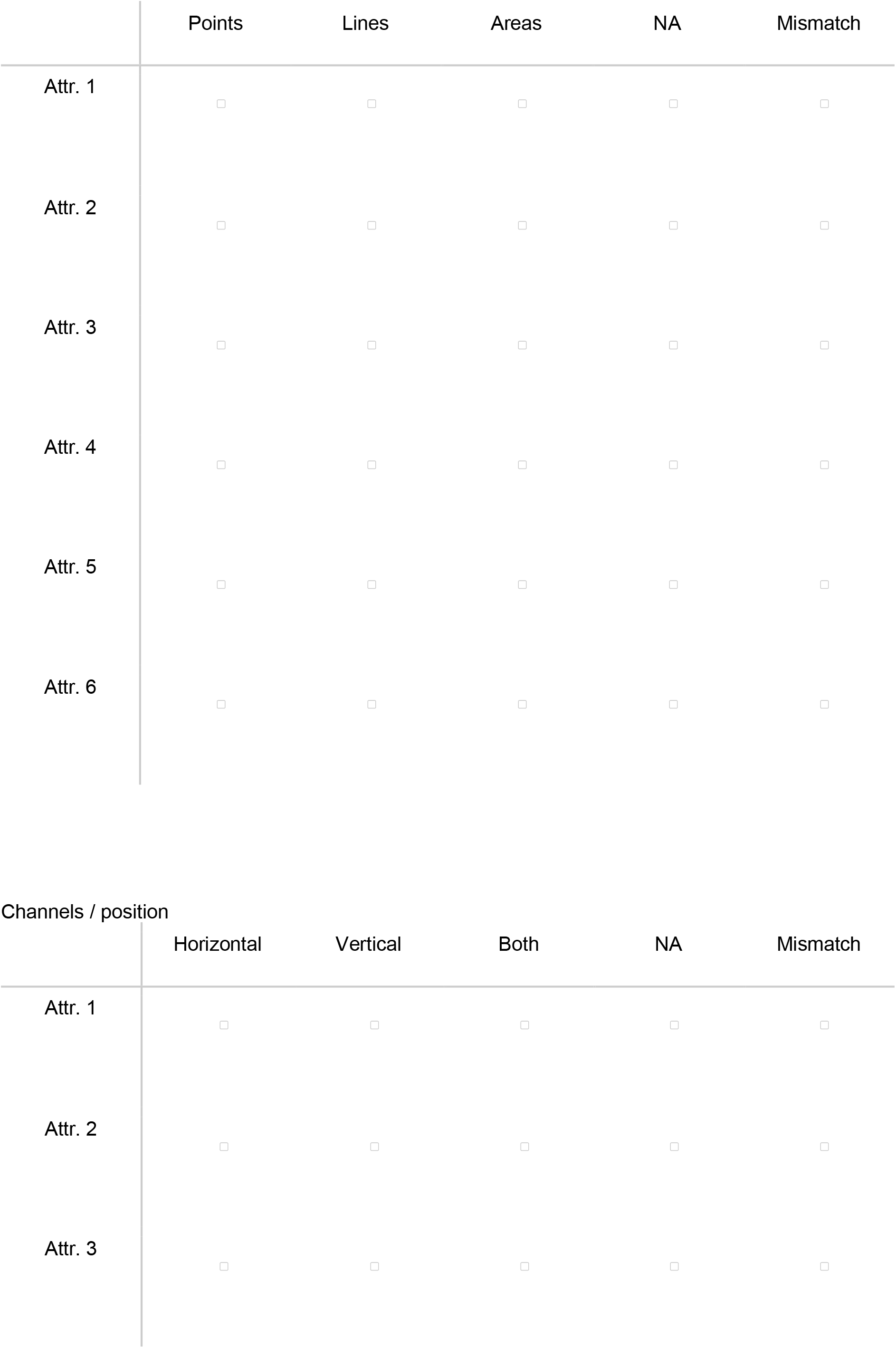

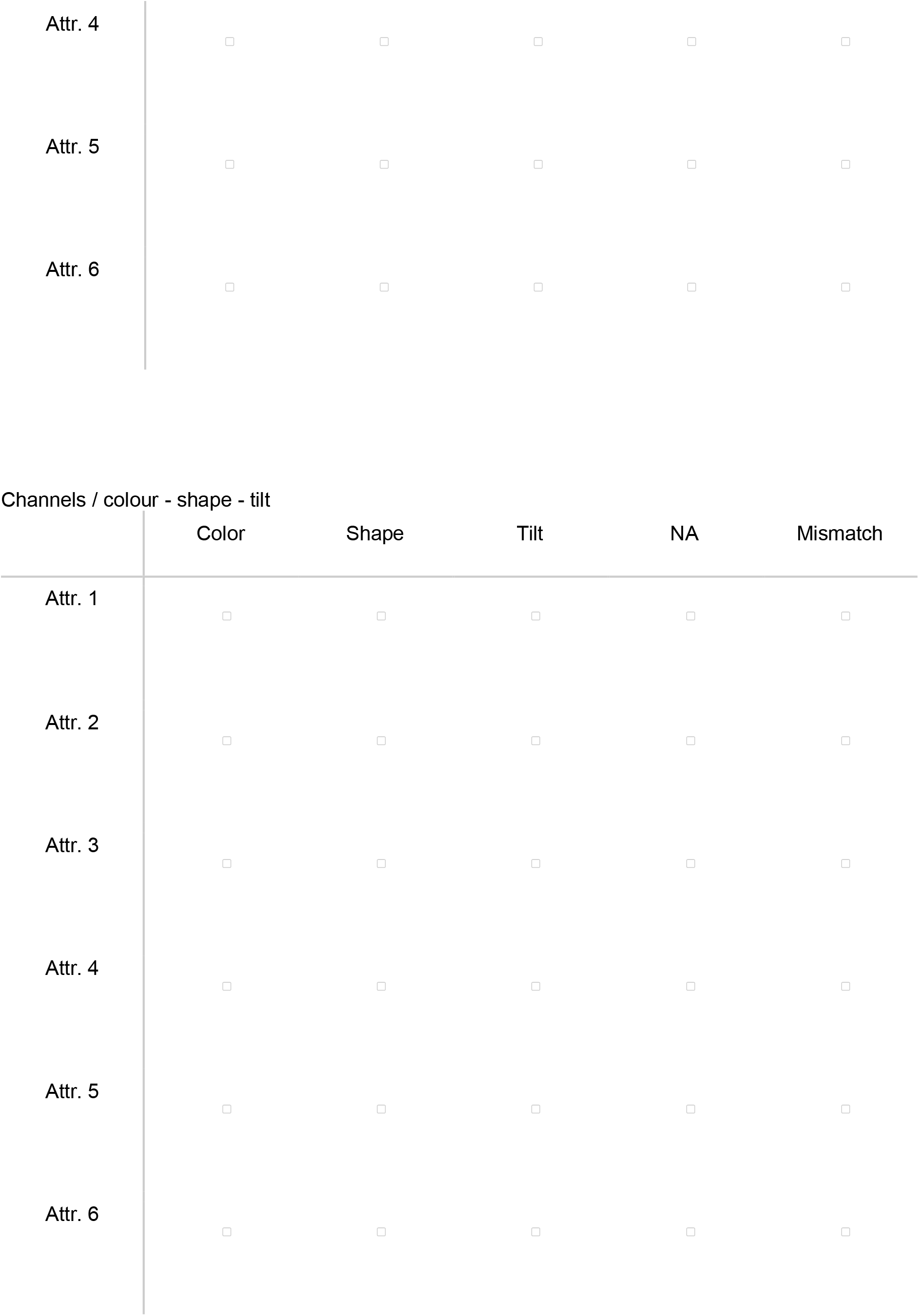

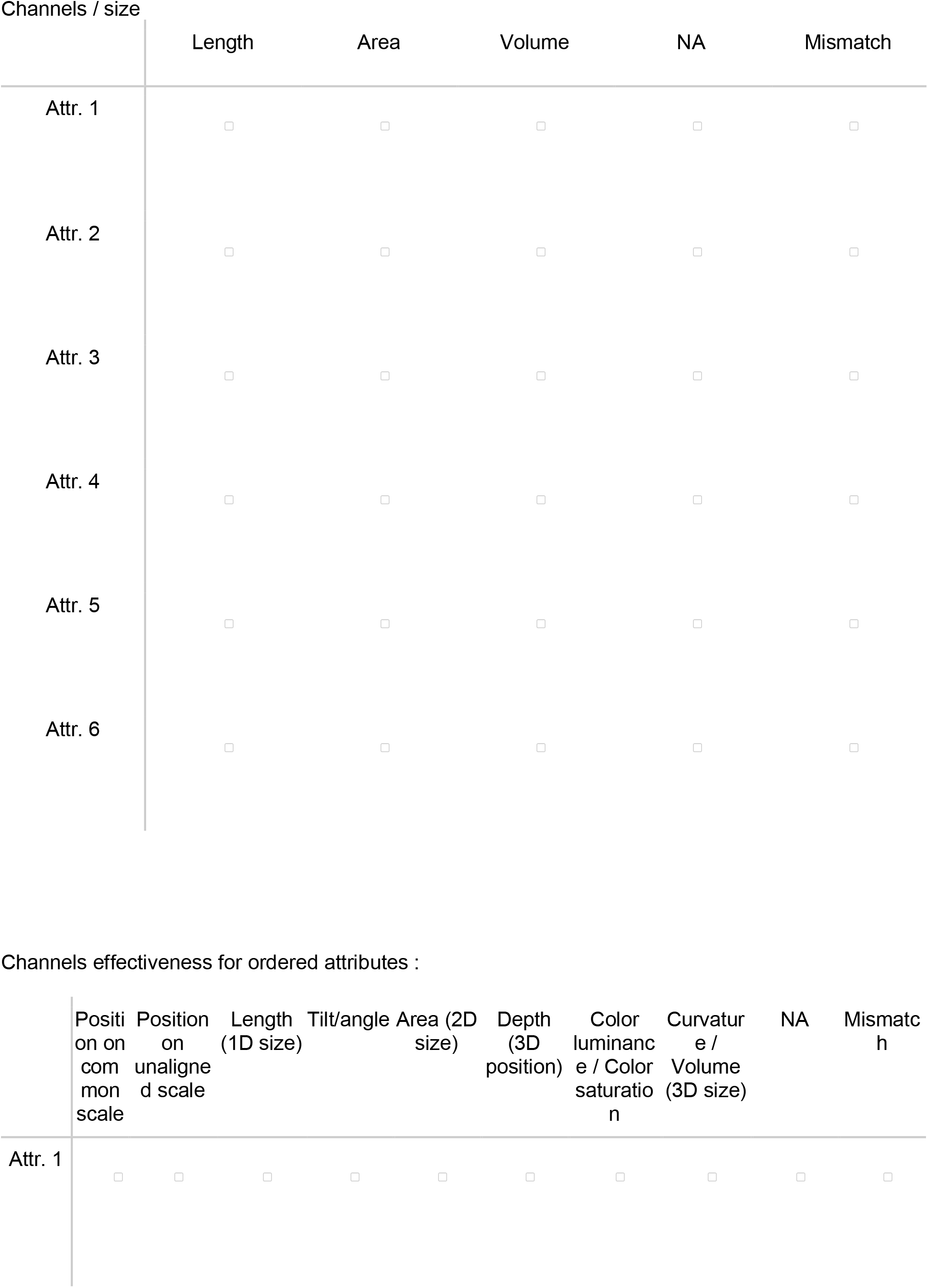

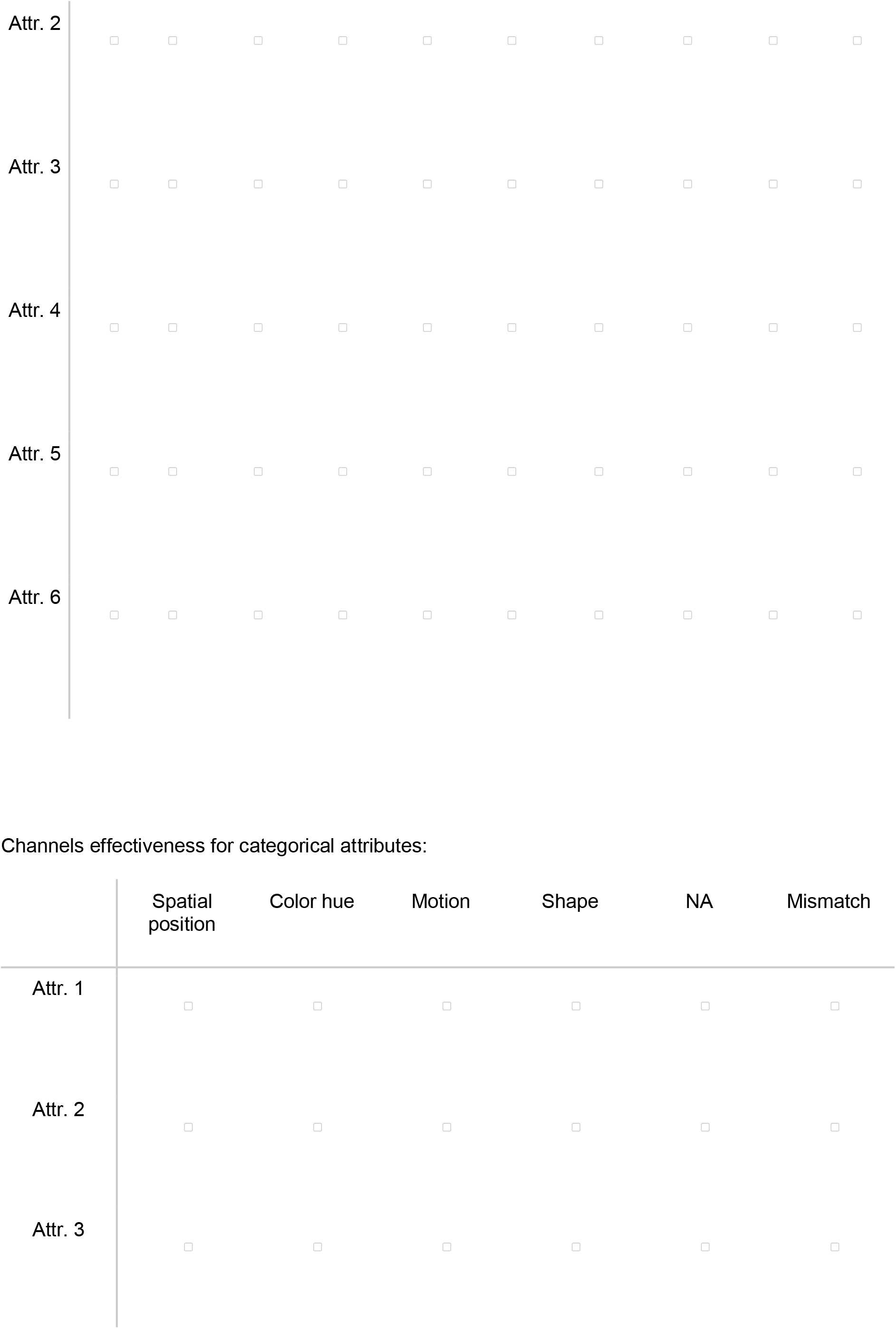

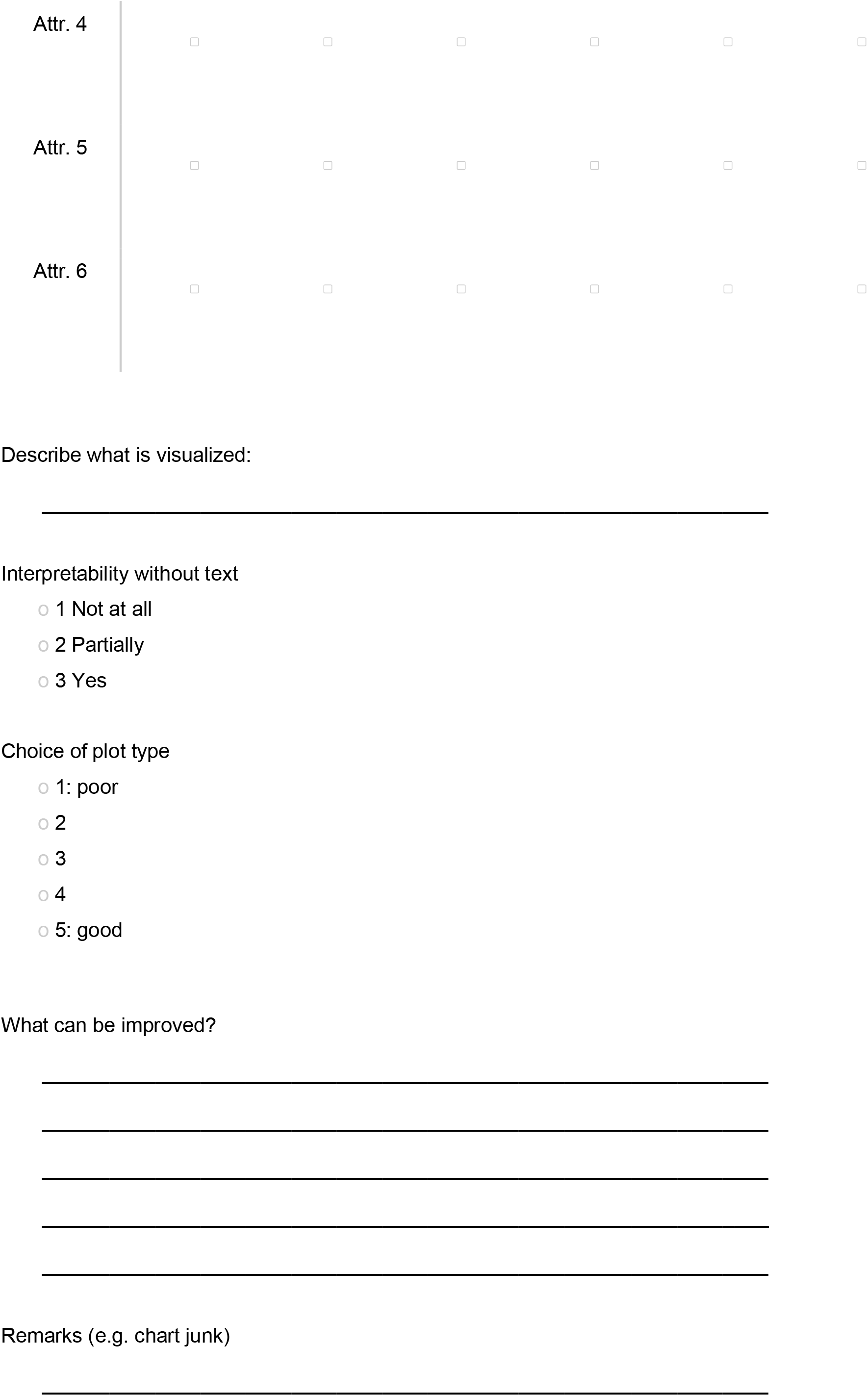

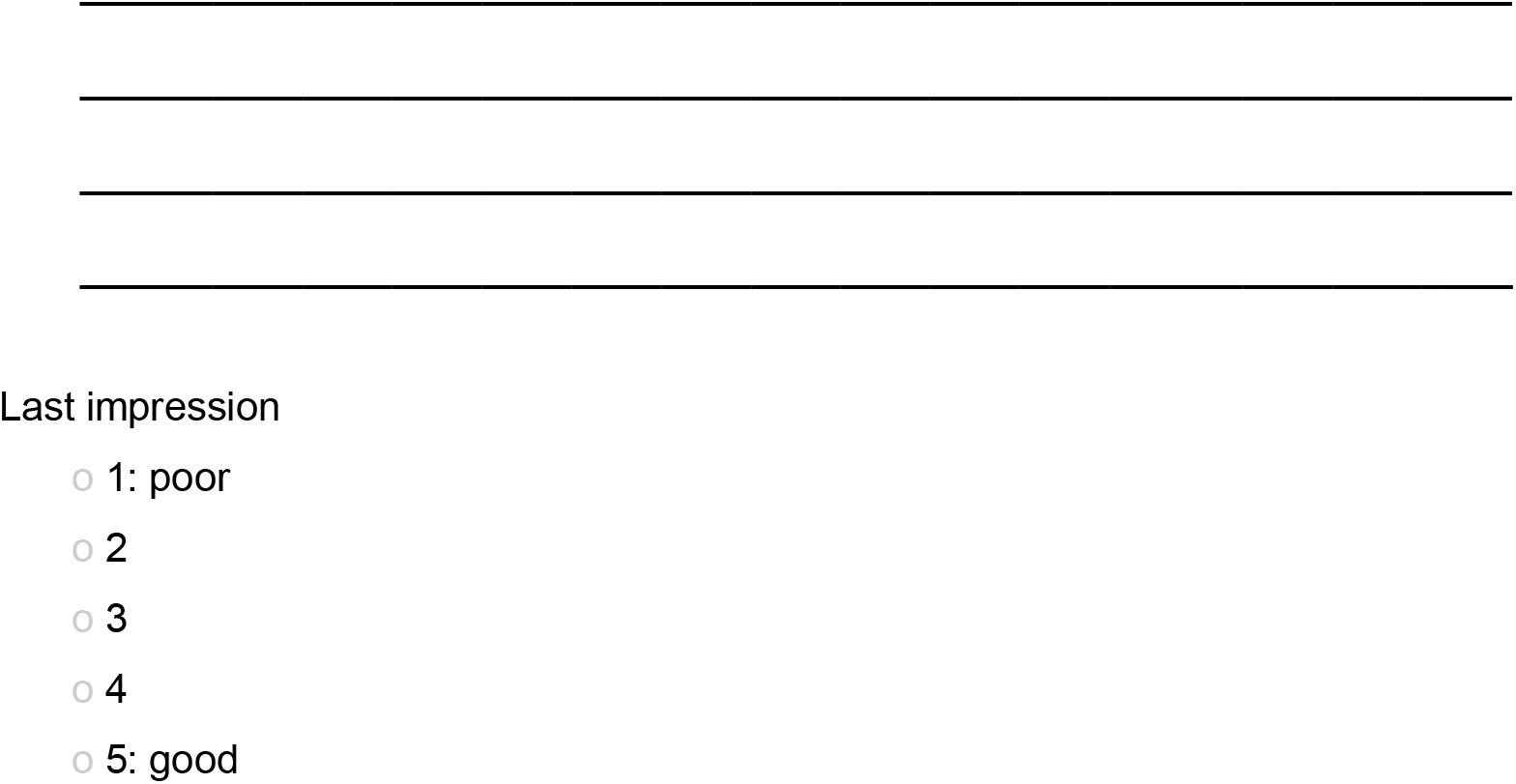

### S2. Visualization types per attribute

**Table S2.**
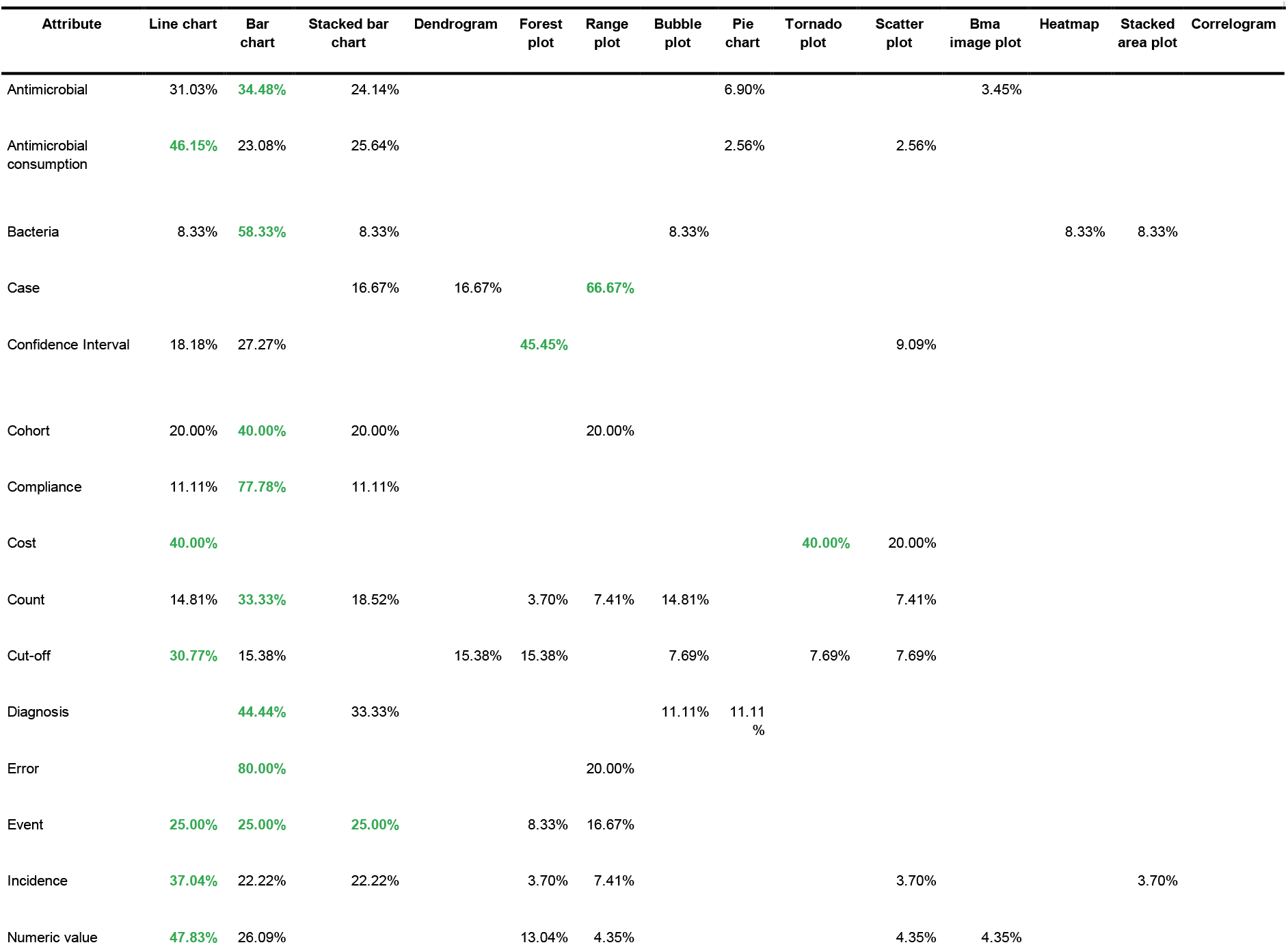

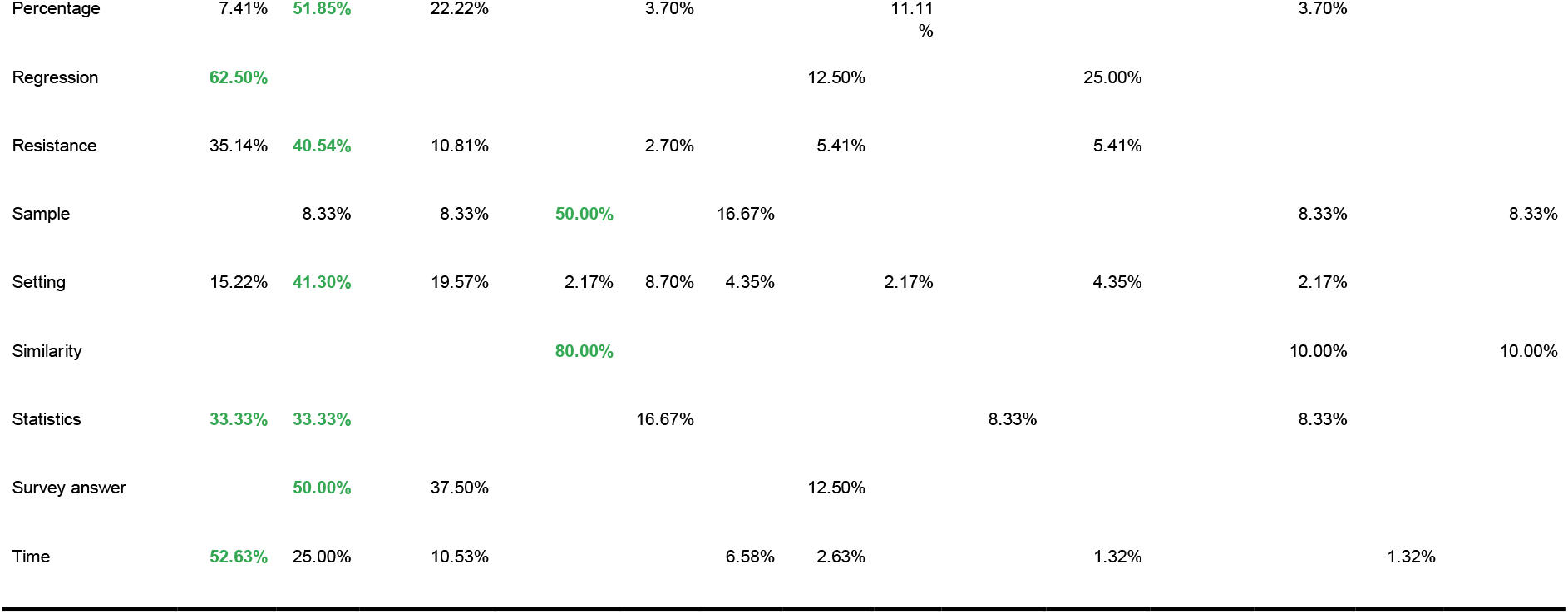

### S3. Visual characteristics per attribute

**Table S3.**

| Attribute | Channel - position |  |  | Channel - marks |  |  | Channel - colour/shape/tilt |  |  | Channel - size |  |  |
| --- | --- | --- | --- | --- | --- | --- | --- | --- | --- | --- | --- | --- |
|  | Horizontal | Vertical | Both | Areas | Lines | Points | Color | Shape | Tilt | Area | Length | Volume |
| Antimicrobial | 72.73% | 27.27% |  |  |  |  | 64.71% | 35.29% |  |  |  |  |
| Antimicrobial consumption | 3.13% | 96.88% |  | 3.03% | 48.48% | 48.48% | 33.33% | 66.67% |  | 6.25% | 93.75% |  |
| Bacteria | 71.43% | 14.29% | 14.29% |  |  |  | 80.00% | 20.00% |  |  |  |  |
| Case | 20.00% | 80.00% |  |  |  |  |  |  |  |  |  |  |
| Confidence interval | 60.00% | 30.00% | 10.00% |  | 81.82% | 18.18% |  | 100.00% |  |  | 100.00% |  |
| Cohort | 40.00% | 60.00% |  |  |  | 100.00% | 75.00% | 25.00% |  |  |  |  |
| Compliance |  | 100.00% |  |  | 83.33% | 16.67% | 100.00% |  |  |  | 100.00% |  |
| Cost | 40.00% | 60.00% |  |  | 50.00% | 50.00% |  |  |  |  | 100.00% |  |
| Count | 23.81% | 76.19% |  | 5.26% | 63.16% | 31.58% | 100.00% |  |  | 18.75% | 75.00% | 6.25% |
| Cut-off | 66.67% | 11.11% | 22.22% | 12.50% | 75.00% | 12.50% | 100.00% |  |  | 50.00% | 50.00% |  |
| Diagnosis | 80.00% | 20.00% |  |  |  |  | 75.00% | 25.00% |  |  |  |  |
| Error | 40.00% | 60.00% |  |  | 100.00% |  | 100.00% |  |  |  | 100.00% |  |
| Event | 100.00% |  |  |  | 33.33% | 66.67% | 25.00% | 75.00% |  |  | 100.00% |  |
| Incidence | 8.70% | 91.30% |  | 5.00% | 45.00% | 50.00% | 28.57% | 71.43% |  | 9.09% | 90.91% |  |
| Numeric value | 55.00% | 40.00% | 5.00% | 6.67% | 40.00% | 53.33% |  |  |  | 33.33% | 66.67% |  |
| Percentage | 34.78% | 60.87% | 4.35% | 12.50% | 75.00% | 12.50% | 66.67% | 33.33% |  | 8.70% | 78.26% | 13.04% |
| Regression |  | 71.43% | 28.57% |  | 71.43% | 28.57% | 33.33% | 66.67% |  |  |  |  |
| Resistance | 7.41% | 92.59% |  | 3.70% | 51.85% | 44.44% | 77.78% | 22.22% |  | 7.14% | 92.86% |  |
| Sample |  | 100.00% |  |  |  |  | 100.00% |  |  |  |  |  |
| Setting | 60.87% | 39.13% |  |  | 33.33% | 66.67% | 59.38% | 40.63% |  |  |  |  |
| Similarity | 100.00% |  |  |  | 100.00% |  |  |  |  |  | 100.00% |  |
| Statistics | 22.22% | 77.78% |  |  | 50.00% | 50.00% | 66.67% | 33.33% |  |  | 100.00% |  |
| Survey answer | 40.00% | 60.00% |  |  |  | 100.00% | 75.00% | 25.00% |  |  |  |  |
| Time | 100.00% |  |  |  | 100.00% |  | 55.56% | 33.33% | 11.11% |  | 100.00% |  |

### S4. Channel effectiveness for quantitative, ordinal and categorical attributes

**Table S4.**
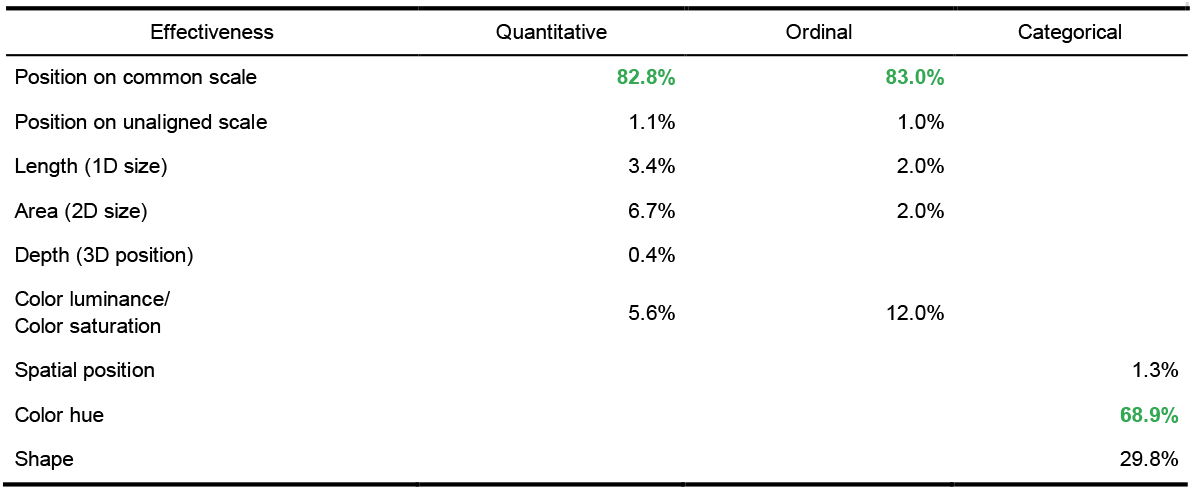

### S5. Problems and illustrative quotes

**Table S5.**
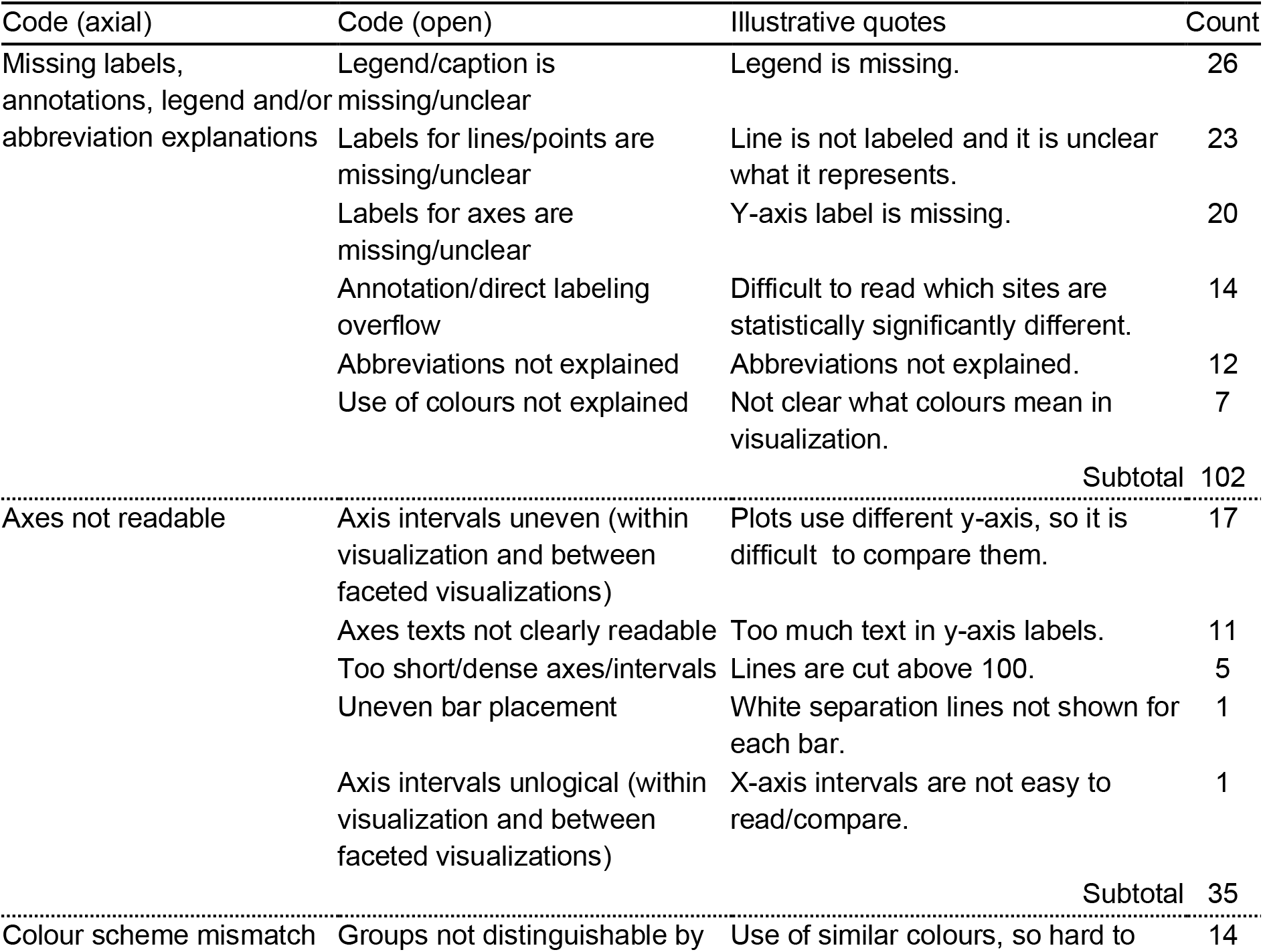
Problems and illustrative quotes

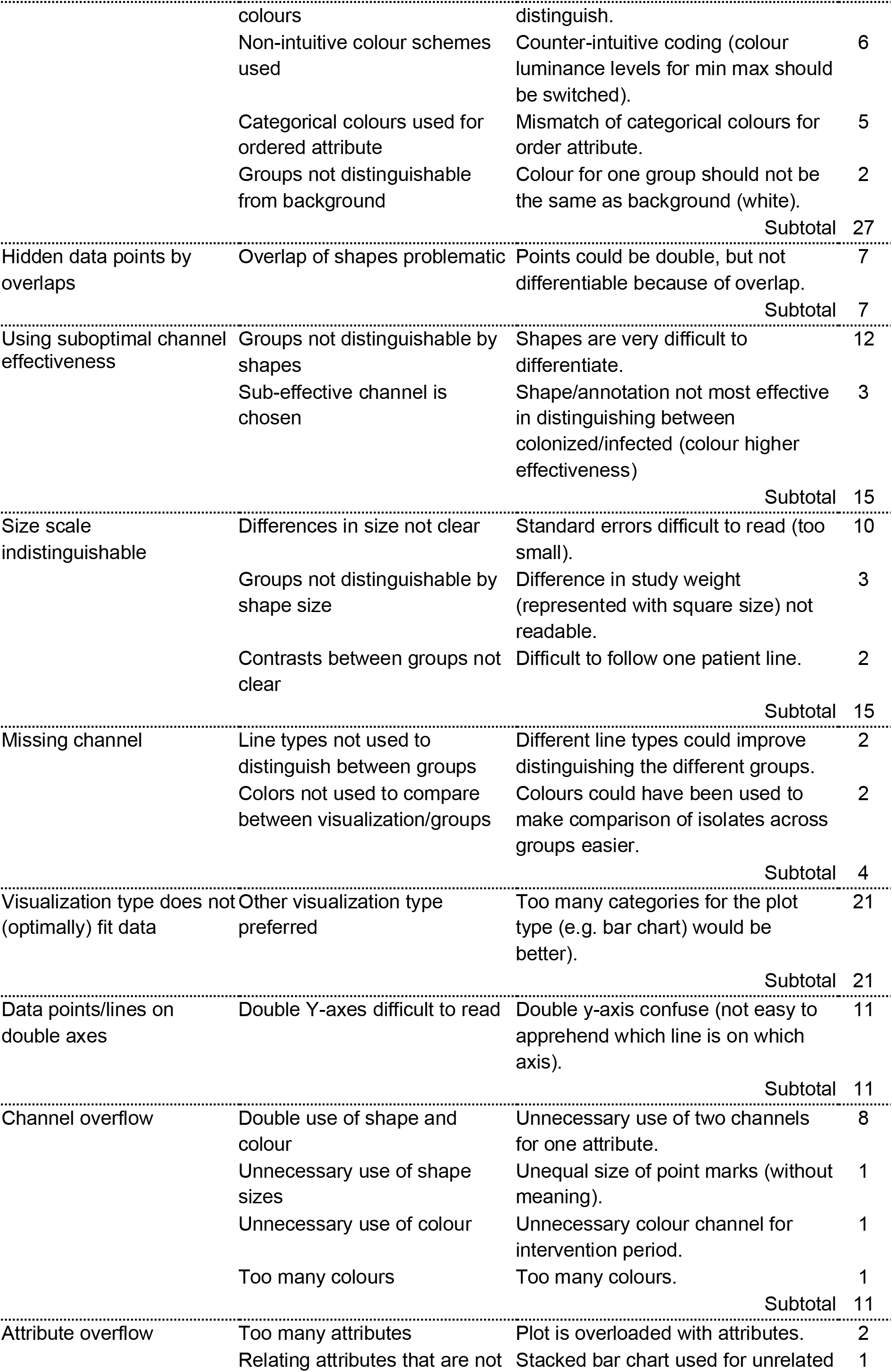

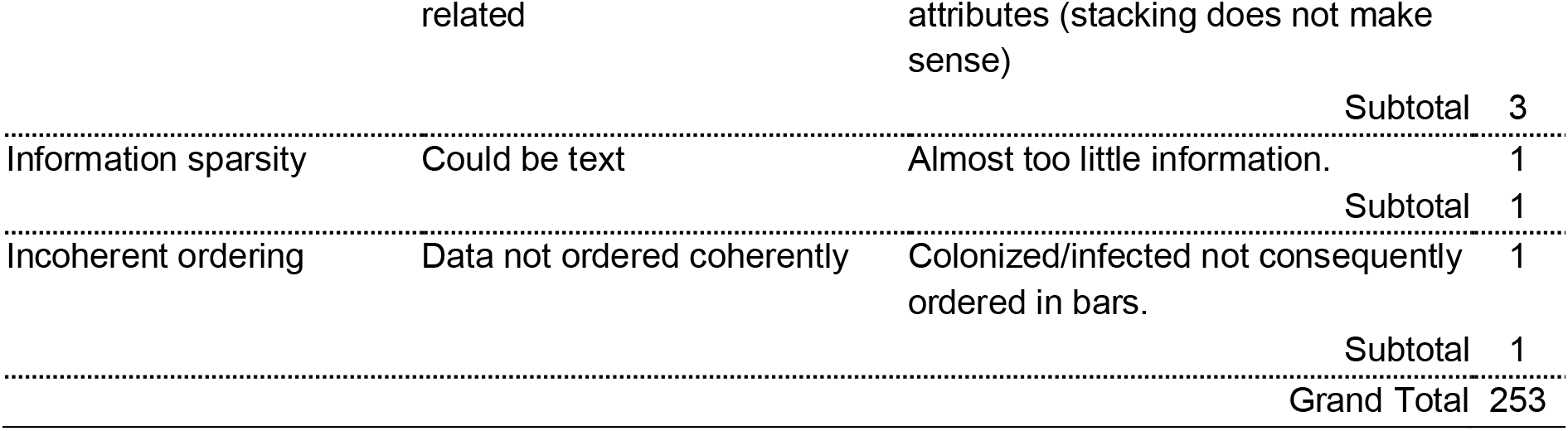

